# THEOBROMA: an aggregated open database of 1.13 million natural products with per-compound license auditing, three-tier classification, and stereochemistry-aware deduplication

**DOI:** 10.64898/2026.06.12.731585

**Authors:** Thor Klamt, Anne Jaczkowski, Jakob Franke, Wolfgang Nejdl

## Abstract

Natural products remain one of the most productive sources of pharmacologically active compounds for drug discovery. Open aggregators attribute licenses at database granularity, and the consequences of that choice have become tangible as the field grows. A recent relicensing event in one constituent source (the September 2024 transition of the Natural Products Atlas to CC BY-NC 4.0) demonstrates how database-level licensing propagates across an aggregate and motivates the per-compound audit framework presented here. The same peer cohort separately leaves classification provenance and stereoisomer-family relations coarser than either layer warrants. THEOBROMA, accessible at https://theobroma.l3s.uni-hannover.de, integrates 1,132,805 natural products from 29 open sources under a per-compound license audit that resolves each compound’s license tier across all attesting sources under a most-restrictive-wins rule, identifying 900,103 compounds (79.5%) under open-use licenses and providing the per-source attestation chain and resolved tier through a dedicated audit endpoint and a query-time license filter. A three-tier classification labels all but 21 compounds by the provenance of the assignment (55.3% curated source labels, 18.2% NPClassifier tool assignments, 26.5% model-inferred). Deduplication at the full 27-character InChIKey retains 1,132,805 entries across 486,032 connectivity families, exposed via a dedicated /api/stereoisomers/<COMP id> endpoint and a radial-family display. Per-compound license provenance is the primary differentiator, while classification stratification and stereoisomer-family exposure add finer-grained access to two related axes, supporting license-compatible virtual screening and isomer-specific bioactivity analysis at corpus scale. As an evolving open resource, THEOBROMA pairs continuous pipeline maintenance with interactive geographic, taxonomic, and chemical-space exploration.

**Graphical Abstract:** THEOBROMA integrates 1,132,805 natural products from 29 open sources across three axes. A per-compound license audit resolves each compound to the most restrictive tier attested by any source, giving 891,860 permissive open, 225,536 non-commercial, 8,243 public domain and 7,166 unspecified, so that 900,103 compounds (79.5%) are open-licensed. A three-tier classification records the provenance of each label as source-supplied (55.3%), tool-assigned (18.2%), or model-inferred (26.5%), leaving 21 compounds unclassified. Deduplication at the full 27-character InChIKey retains the stereochemical and protonation variants that 14-character truncation would erase, illustrated by the (E,E)- and (Z,E)-curcumin pair shown.

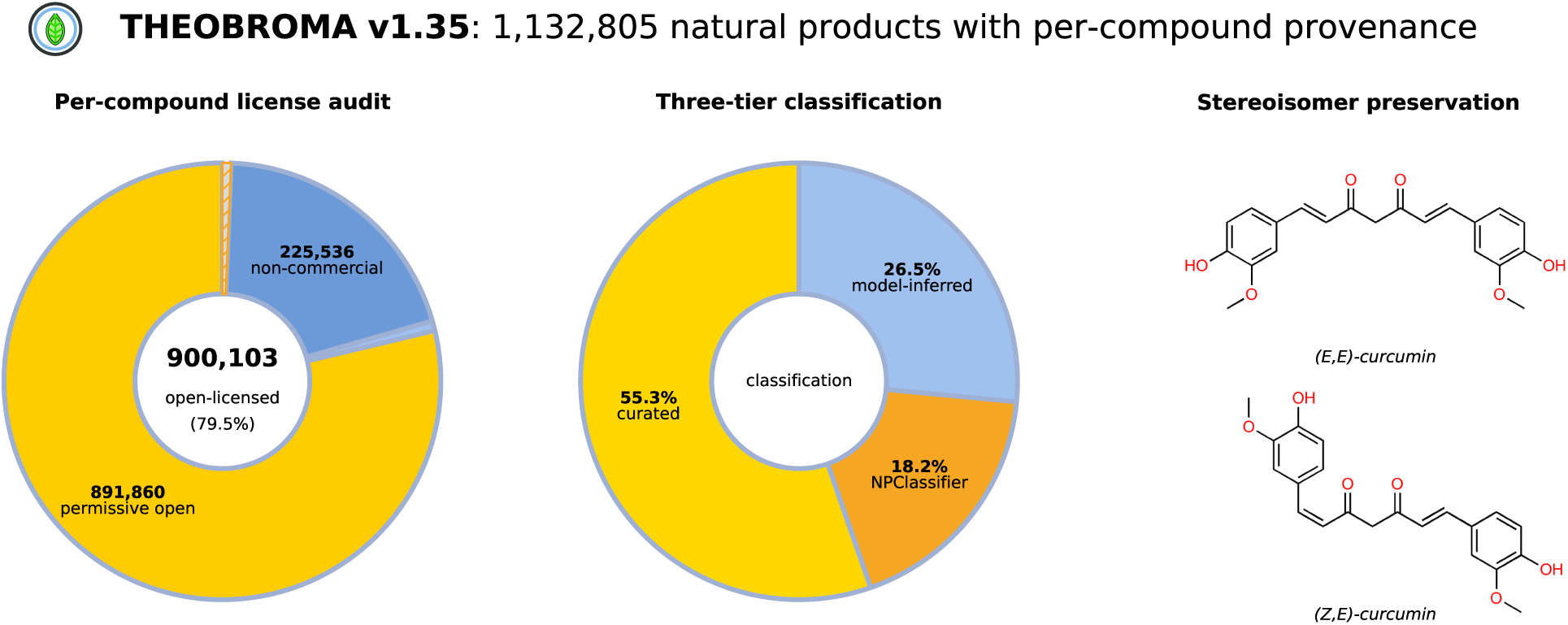

## Introduction

Natural products account for roughly one-third of small-molecule drugs approved between 1981 and 2019 as direct derivatives and close to two-thirds when scaffold-inspired compounds are included (1; 2). The September 2024 transition of the Natural Products Atlas from CC BY 4.0 to CC BY-NC 4.0, a permitted exercise of the maintainers’ rights following commercial repackaging of the open corpus into the AntiBase product, illustrates the structural fragility of database-level license attribution in the open natural-product aggregator ecosystem: a relicensing event at any single source propagates across the entire aggregate, removing commercial reuse even for compounds that did not originate from the relicensed source (3). Current open aggregators group broadly into three overlapping emphases. Scale-oriented resources contribute the bulk of the catalogued chemistry, with COCONUT 2.0 integrating 63 source collections at 695,133 unique structures under a stereochemistry-aware data model and LOTUS hosting roughly 750,000 structure-organism pairs on Wikidata, with a Zenodo export under separate terms (Supplementary S3) (4; 5). The remaining resources are curation-oriented or domain-focused. Curation-oriented and domain-focused resources cover the Natural Products Atlas 3.0 for microbial compounds, NPASS 3.0 for quantitative composition and experimental ADME-Tox records, HERB 2.0 for traditional Chinese medicine, IMPPAT 2.0 for Indian medicinal plants with an explicit stereochemistry-aware library, and SuperNatural 3.0 for vendor-available compounds with a three-level annotation confidence indicator (3; 6; 7; 8; 9). The peer cohort has converged on shared conventions: license attribution at database granularity, aggregate classification reporting without provenance stratification, and in the most recent releases stereochemistry-preserving deduplication.

Three problems remain unresolved against this background. License attribution at database granularity proves fragile under commercial-use pressure. PubChem displays per-source license information within individual compound records, so per-source metadata at compound granularity is not new. The audit adds the resolution step: heterogeneous attestations are collapsed into a single tier under a stated precedence order, materialized as a queryable column and exposed through an audit endpoint. Aggregate classification coverage, reported as a single collapsed figure, conflates source-supplied labels with model-inferred ones at heterogeneous confidence. Here too the idea is not new. UniProtKB has long separated reviewed from unreviewed entries and carries a five-level annotation score (10). The distinction here is that stratification is applied within the classification axis and tied to the provenance of each assignment, ordered as curator, tool, and model, where COCONUT 2.0 and SuperNatural 3.0 aggregate across all annotation axes into a single per-compound quality score (4; 9). Stereoisomer preservation is now the norm, with COCONUT 2.0 retaining stereochemically defined molecules as distinct entries and IMPPAT 2.0 offering an explicit stereochemistry-aware library (4; 8). The remaining gap is exposure: the family relation is retained by several peers but not offered as a directly queryable resource.

THEOBROMA v1.35 addresses these gaps. Against database-level attribution, the licensing audit carries the terms of every attesting source through to each compound and resolves them conservatively, identifying 900,103 compounds (79.5%) under open-use licenses. A relicensing event is contained to the compounds its source attests and does not propagate across the aggregate. The three-tier classification records who assigned each label, not how confident the assignment is, keeping curator, tool and model annotations separable at query time. Stereoisomer families are reachable through their own endpoint and rendered per compound. The corpus is 1,132,805 compounds from 29 open sources spanning four biological kingdoms plus an unresolved bucket, audited in March 2026 and accessible at https://theobroma.l3s.uni-hannover.de. Geographic provenance, traditional-medicine tagging, 47 in silico annotation columns and a FAIR-compliant (11) reproducibility manifest are described below, and Table 1 summarizes the differentiator axes against the peer cohort.

**Table 1.**
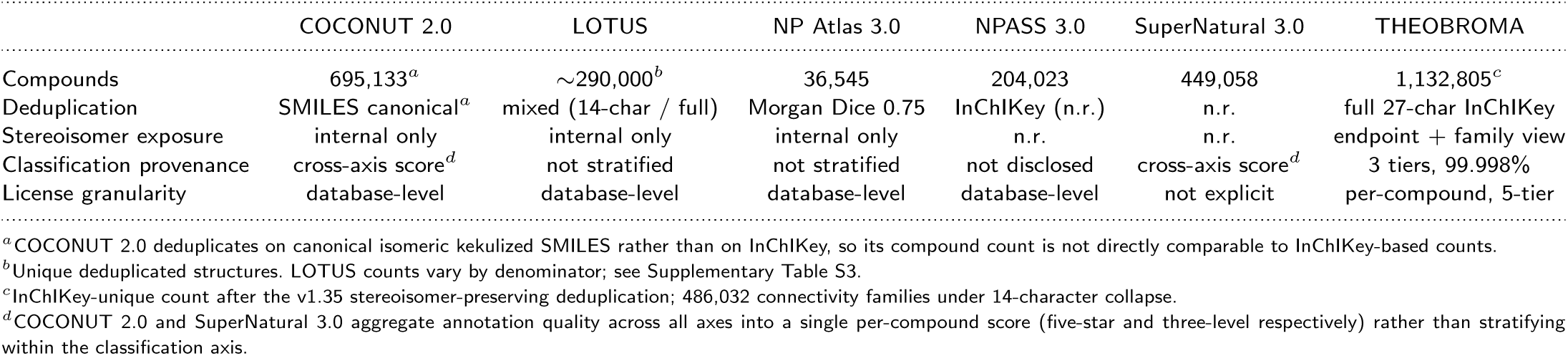
THEOBROMA against the five most directly relevant open natural-product aggregators, restricted to the three differentiator axes. Source counts, kingdom coverage, similarity search, geographic annotation, and ADMET content are compared in Supplementary Table S3. “n.r.” abbreviates “not reported”.

### Data Sources and Integration

THEOBROMA aggregates 29 sources spanning six overlapping emphases: large-scale aggregators dominate at 998,699 compounds (COCONUT 2.0, LOTUS, NPASS 3.0, Natural Products Atlas), with food and metabolomics resources contributing 58,775 (FooDB, LMDB Lichen, CMDB Cereals) and traditional-medicine, regional and geographic, microbial and fungal, and phytochemistry-focused resources contributing the remaining 75,331. Supplementary Table S4 lists per-source counts, versions, and licenses. Each source is documented in a YAML reproducibility manifest recording download date, version, license, citation, source URL, and the integration script applied during ingestion (Figure 1).

**Figure 1.**
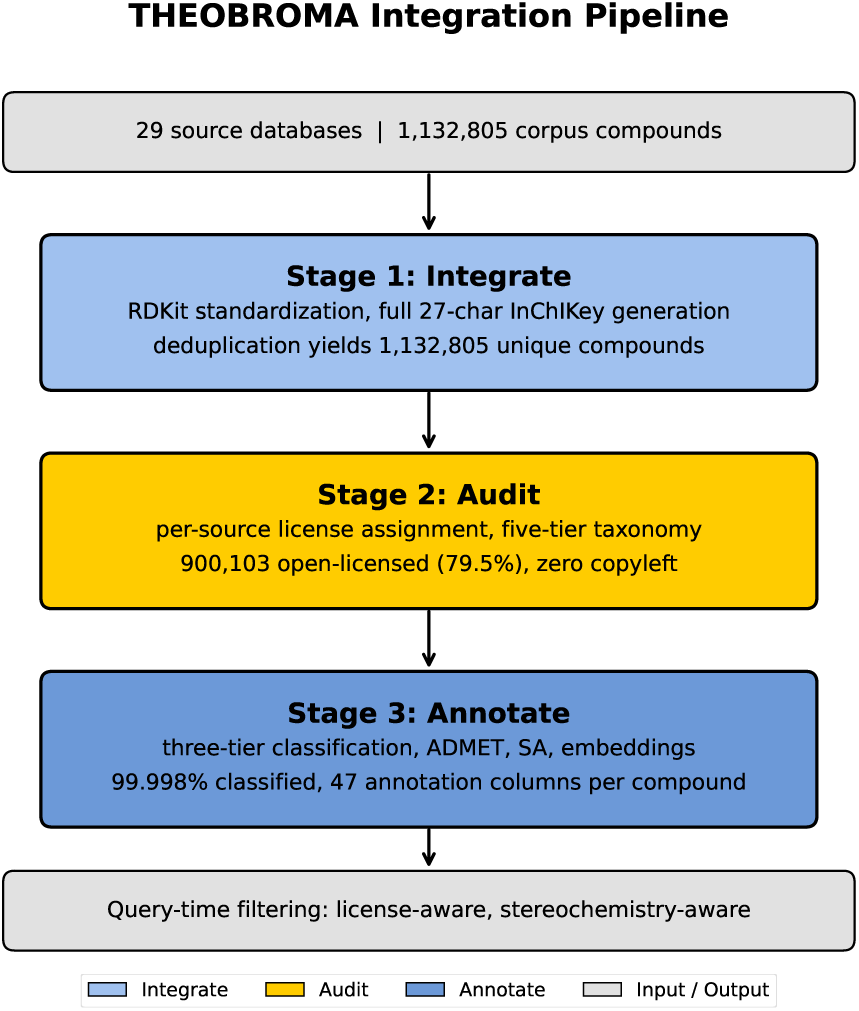
THEOBROMA integration pipeline. Three stages carry 29 source databases into the v1.35 corpus of 1,132,805 compounds: Integrate applies RDKit standardization and full 27-character InChIKey deduplication, Audit assigns a license tier per source and resolves each compound under a most-restrictive-wins rule, and Annotate adds the three-tier classification, 47 in silico annotation columns, and learned embeddings. The output banner shows the license-aware and stereochemistry-aware filtering the pipeline enables.

Compound provenance is resolved through a three-authority taxonomy combining the World Checklist of Vascular Plants (12) for plant lineages, the NCBI Taxonomy database (13) for microbial and animal lineages, and the World Register of Marine Species (14) for marine organism cross-validation, materialized in a canonical resolved taxonomy table that assigns each compound a primary kingdom and an optional secondary kingdoms array. Marine organisms are assigned by lineage rather than treated as a kingdom, so marine-derived compounds resolve to animal, fungi, plant, bacteria, or unresolved. Protist-supergroup compounds are assigned to unresolved rather than forced into one of the four kingdoms. The distribution is dominated by plant (964,360), animal (80,485), and fungi (41,545), with bacteria (32,749) and unresolved (13,666) completing the corpus.

The resolution also infers kingdom from phylum-level WCVP attestations. Primary kingdom is assigned by majority vote across a compound’s organism attestations. An attestation resolving to the phylum Streptophyta but carrying no kingdom is inferred as plant and included in that vote, recovering the dominant case of NCBI- resolution bias that would otherwise exclude WCVP-resolved plants from the count. The secondary kingdoms array in resolved taxonomy carries all attested kingdoms regardless of resolution status, and users querying compounds with multi-kingdom attestation should consult the array alongside the primary kingdom assignment. Smaller residual cases (Rhodophyta, Evosea, Oomycota, Euglenozoa) and finer source-weighting refinements remain open for a future release.

Standardization is applied uniformly across sources using RDKit (15), as detailed in Supplementary S1, and salts are preserved as multi-fragment SMILES (5,443 entries, 0.5%). Against that principle, 199 trivially small species carrying three or fewer heavy atoms are removed, 49 of them metal-bearing, since they carry no scaffold, no stereochemistry, and no biosynthetic classification. Removal precedes deduplication and per-source attribution, so the counts of Supplementary Table S4 are taken after it.

Deduplication operates on the full 27-character InChIKey (16), producing 1,132,805 unique entries in 486,032 stereoisomer-and- protonation families. Family membership is defined by the 14- character connectivity prefix. Stereochemistry, isotopic substitution, and protonation state are encoded in the later blocks, so family members share connectivity while differing in one or more of those states.

Ingestion is sequential. COCONUT 2.0 is processed first and the remaining 28 sources are appended by full-InChIKey matching, so new compounds extend the corpus while overlapping compounds aggregate metadata into the existing entry, which makes primary attribution a function of ingestion order rather than of source precedence, yielding 481,817 synonyms across 149,178 compounds (3.23 per named compound on average) and the per-compound source-license, pathway, and kingdom attributions that ground the audit of the following section. The 919,873 compounds attributed to COCONUT therefore exceed the 695,133 of its own publication by 224,740, reflecting stereoisomer, protonation, and isotopologue variants that its coarser convention collapses. Collapsing protonation while retaining stereochemistry, the convention that figure follows, gives 808,579 entries. Collapsing stereochemistry as well gives 486,032. Of the 458,432 entries (40.5%) carrying a non-neutral protonation flag, 134,206 are the only entry for their skeleton and stereochemical configuration, so collapsing protonation would remove them rather than merging them.

### Licensing Audit

The Natural Products Atlas relicensing event introduced above motivates the per-compound audit described here. Downstream users of the post-September-2024 Atlas face a binary choice between accepting CC BY-NC 4.0 across all 36,545 compounds in that corpus or manually re-auditing each contributing source to reconstruct the unaffected fraction. That figure is the Atlas corpus in full, of which 5,513 compounds enter THEOBROMA under primary attribution (Supplementary Table S4). THEOBROMA’s per-compound audit, applied across its 29 integrated sources, removes this re-audit burden by providing source-level licenses at compound granularity at query time.

The audit was performed per source in March 2026. License metadata was extracted in order of precedence from upstream repositories, peer-reviewed citations, download or terms-of-service pages, and direct correspondence with maintainers. Each source was then assigned to one of five tiers: permissive open, public domain, non-commercial, copyleft, and unspecified (Supplementary S3). Conservative reattribution was applied where data-level licensing was ambiguous rather than absent, moving four sources from permissive open to non-commercial. Share-alike sources were excluded. CMNPD (CC BY-NC-SA 4.0) falls outside the commercially reusable fraction on its NonCommercial term alone, and its ShareAlike term would additionally propagate to the aggregate if the integration were treated as an adaptation rather than a collection (Supplementary S3).

A second stage resolves each compound’s license tier across all attesting sources under a most-restrictive-wins rule. Any compound attested by a non-commercial source resolves to non-commercial regardless of permissive co-attestations, any compound attested by an unspecified-license source resolves no more permissively than unspecified, and co-attestation by CC BY 4.0 and CC0 sources resolves to CC BY 4.0, since the attribution requirement must be honoured downstream. The resolved tier is materialized in the license tier column of the compounds table, and the per-source attestation chain that supports the resolution is persisted in a dedicated per source license attestation table and exposed via a /api/compound/ <ID>/license-provenance endpoint so that downstream users can audit the resolution for any individual compound. The resolver is applied as a single SQL transaction with idempotent behaviour and is documented in Supplementary S3. The rule is a conservative operational choice rather than a legal determination: honoring the most restrictive attestation is the interpretation under which no attesting source’s terms can be breached. The audit records the terms each maintainer asserts over the data it distributes and does not adjudicate whether those terms are enforceable over an individual record, a question that depends on jurisdiction and on whether copyright, the sui generis database right, or contract is the operative instrument (Supplementary S3). The resolved tier is a compliance floor, not legal advice.

Applied across the corpus, the resolver places 891,860 compounds (78.7%) under permissive open licenses and 8,243 (0.7%) under public domain, giving an open-licensed fraction of 900,103 (79.5%). The remainder resolves to non-commercial (225,536) or unspecified (7,166). No compound resolves to copyleft (Supplementary Figure S1). Per-source assignments are given in Supplementary Table S4, whose notes explain why the per-source rows do not sum to these totals. Commercial users assemble license-compatible subsets through the query-time license filter.

The resolver also records the least restrictive tier attested for each compound. The two bounds answer different questions. The most restrictive tier is the license under which a compound may be redistributed as part of the aggregate, honoring every attesting source simultaneously, and is the default reported throughout. The least restrictive tier is the best license under which the compound is attested by any single source, and is therefore a sourcing figure rather than a redistribution one: a user who obtains the compound from that source directly, and attributes it accordingly, is bound by that source’s terms alone. This matters for 137,149 compounds (12.1%). Each carries attestations spanning more than one tier, so the resolved tier understates what is obtainable by approaching a single source. This exceeds the 103,304 compounds that the resolver moves to a more restrictive tier (Supplementary Table S4), since for the remaining 33,845 the primary source already carried the most restrictive attestation. Under the least restrictive bound the permissive-open fraction is 1,014,300 compounds and the public-domain fraction 18,925, so 1,033,225 compounds (91.2%) are attested under CC BY 4.0 or better by at least one source. The resolved tier reported throughout, and used by the query-time license filter, is the most restrictive. The least restrictive bound is exposed per compound through the audit endpoint. Both bounds depend on a single source at the public-domain end. TM-MC is the only CC0 attestation in the corpus. Under the least restrictive rule all 18,925 compounds it attests reach public domain. Under the most restrictive only the 8,243 it attests alone do, since any co-attestation carries an attribution requirement. A relicensing at that source would eliminate the tier under either. The same fragility applies at far greater scale at the permissive-open end, where COCONUT’s CC BY 4.0 designation carries 919,873 compounds under primary attribution and therefore underwrites most of the open-licensed fraction. This is the single-source fragility the per-compound audit is designed to contain: the audit does not prevent a relicensing, but records which compounds are affected, so the rest remain available under their own terms.

### Chemical Descriptors and Annotations

Each compound carries four annotation layers covering classification, chemical descriptors and embeddings, traditional-medicine provenance, and similarity-based target prioritization, with per-layer methodology in the subsections that follow and full detail in Supplementary S4 to S6, S9 and S10.

#### Three-tier classification

Class assignments carried in the source databases are accepted directly for 626,601 compounds (55.3%, curated tier). Compounds without a curated label are assigned by the NPClassifier tool itself where it returns a prediction (206,042 compounds, 18.2%, tool tier). The remainder are assigned by an XGBoost ensemble (17) over the NPClassifier class vocabulary, operating on a fingerprint plus embedding feature vector with hyperparameters selected via Optuna (18) (300,141 compounds, 26.5%, model tier, Supplementary S4). Twenty-one compounds remain unclassified because no feature vector could be computed for them. The curated tier records that the label was supplied by the source, not that a human assigned it. Several sources, COCONUT among them, generate their own NPClassifier or ClassyFire annotations (Supplementary S4). The curated tier is larger than in v1.34 (626,601 against 397,278) since the tier assignment is materialized from source provenance in this release, where the preceding version derived it from model confidence. The two figures are therefore not measurements of the same quantity, and no new curated labels were acquired from upstream. The tier ordering records which source assigned the label, not a reliability score, and the per-compound model confidence is exposed alongside the assignment so that downstream users can impose their own acceptance threshold without re-deriving the classification. Superclass and pathway are derived from the class through the NPClassifier ontology under a designated primary parent, so all three tiers share a single vocabulary at every rank. Five-fold cross-validated macro-F1 is 0.809 *±* 0.002 and held-out macro-F1 on the 57,601-compound partition is 0.825. The deployed model is refitted on the full training partition rather than on the four-fifths available to each fold, and the held-out figure exceeds the cross-validated mean for that reason. Per-class precision-recall structure and confusion matrices are provided in the project repository, and the search-space and selected-value detail is in Supplementary S4.

Pathway is reported from effective pathway, derived from the assigned class rather than inherited from the source, and covers 1,132,653 compounds. The largest category is “Alkaloids” at 26.9%, narrowly ahead of “Terpenoids” at 24.2% and “Shikimates and Phenylpropanoids” at 18.2% (Supplementary Figure S3). The category names are reported as NPClassifier assigns them. Its pathway vocabulary groups compounds by biosynthetic origin, so “Alkaloids” denotes nitrogen-containing compounds of diverse biosynthetic routes rather than a single pathway, and “Shikimates and Phenylpropanoids” denotes shikimate-derived compounds rather than shikimate itself. The source-supplied pathway column is retained but not used, for the reasons given in Supplementary S4.

#### Chemical descriptors and embeddings

All but sixteen compounds carry the 41 ADMET-AI endpoint predictions (19), RDKit physicochemical descriptors (15), three structural alert sets (PAINS (20), BRENK (21), and the NIH filter set as distributed with RDKit), and Bemis-Murcko scaffolds (22). Synthetic accessibility scores (23) cover 1,131,184 compounds (99.86%, Supplementary Figure S4) and pathway annotation all but 152. Each compound also carries 768-dimensional mean-pooled embeddings from the original ChemBERTa checkpoint (seyonec/ChemBERTa-zinc-base-v1, pretrained on a 100k-molecule subset of ZINC) (24) indexed via FAISS (25) for sub-linear nearest-neighbor retrieval. ADMET predictions cover a slightly larger 1,132,789-compound subset than the synthetic-accessibility scores, because the two pipelines fail on different molecules, primarily salts and isotopologues. The ADMET regression endpoints are unconstrained model outputs reported as produced, so predictions can fall below the physically meaningful range where the training distribution lies near zero: 354,826 compounds carry a negative predicted volume of distribution and 274,202 a negative predicted half-life. These are model artifacts rather than property estimates. The compound page clamps them to the physical range for display and shows the raw value alongside, while the API and the deposited data carry the model output unchanged. ADMET-AI is trained on Therapeutics Data Commons benchmark sets dominated by drug-like synthetic chemistry, so some fraction of the natural-product corpus lies outside its training distribution. No formal applicability domain is computed, but the novelty axis of the trust score, the normalized Tanimoto distance to the nearest approved drug in DrugBank (Supplementary S10), serves as a first-order proxy: high values indicate distance from the chemistry the models were fitted on. The predictions are best read as a screening-scale annotation layer rather than as property estimates for any individual structure.

Of these, the 41 ADMET-AI endpoints and six RDKit-derived flags are materialized as the 47 columns of the ADMET annotation table, while the physicochemical descriptors sit alongside the structural record. The annotation surface spans these columns together with the classification, taxonomy, geographic and provenance layers, with full methodology in Supplementary S5. A nine-axis trust score quantifies per-compound annotation coverage across that surface at corpus mean 0.326 (Supplementary S10).

#### Traditional-medicine provenance tagging

Traditional-medicine provenance is tagged on 60,242 compounds (5.3%) across Traditional Chinese Medicine and related East Asian systems and Ayurvedic and Indian traditional medicine, of which 4,841 are attested in both (Supplementary Figure S5, with per- tradition counts in Supplementary S6). THEOBROMA inherits the licensing constraints of upstream sources, and users intending commercial development should verify compliance against the originating source’s licensing and any applicable access-and-benefit-sharing frameworks.

A related obligation attaches outside the tagged set: the CSIRO Australian Natural Products collection states that its linked publications may contain Indigenous Cultural and Intellectual Property, and that use of those publications or collection of specimens containing the described compounds may require permission from First Nations peoples. This attaches to the source material rather than to redistribution of the structures, and is therefore orthogonal to the resolved license tier.

#### Similarity-based target prioritization

As a heuristic browsing affordance rather than a predictive feature, per-compound nearest-neighbor similarity is precomputed against 25 reference target panels spanning kinases, GPCRs, nuclear receptors, proteases and ion channels (Supplementary S9). For each compound and each panel, the score is the maximum Morgan Tanimoto coefficient between the compound and the set of known ChEMBL v34 actives for that panel (26; 27). The resulting score vector surfaces candidates structurally similar to known actives, which is useful for triaging a large library, but it says nothing quantitative about whether any individual compound will bind. It is neither a binding-affinity prediction nor an activity classifier nor a substitute for downstream docking or assay-based validation, and both the API response and the compound-detail page label it exploratory for that reason. Users requiring quantitative target-prioritization performance should conduct their own retrospective benchmarking against target-appropriate decoy sets with explicit applicability-domain controls.

### Web Interface and Use Cases

THEOBROMA is served via a Flask web application backed by PostgreSQL, with a REST API documented at /api/docs and a machine-readable OpenAPI specification provided in the Supplementary Materials. No login or registration is required for any search, download, or batch-annotation function, and the web interface is responsive across desktop, tablet, and mobile form factors. Search modes cover textual retrieval across compound names, synonyms, and source identifiers, structure-based retrieval via SMILES with four similarity modes over Morgan and MACCS fingerprints and two learned embeddings, the general-chemistry ChemBERTa representation and the natural-product-pretrained NaFM representation (28), the latter restricted to structures resolvable to a corpus entry, and SMARTS substructure matching. Further modes cover Bemis-Murcko scaffold browsing and a multi-filter advanced query combining structural, classification, ADMET, and provenance constraints. A license-aware filter composes with all of them, resolving against the most restrictive attested tier so that no downstream re-audit is needed. A stereoisomer-family endpoint returns all structures sharing a 14-character InChIKey connectivity prefix with the query, which includes protonation and isotopologue variants alongside stereoisomers. The endpoint name predates the broader family definition adopted here. A batch annotation interface at /annotate accepts structure-list uploads in CSV, TSV, Excel, plain text, or SMILES format with auto-detected SMILES or InChIKey columns, returning per-compound annotation for corpus intersections and an unmatched list for non-corpus structures through chunked client-side calls to the corresponding /api/annotate REST endpoint (Supplementary S12). Median served response time is 40 ms for textual search and under one second for every structure and embedding similarity mode, measured end to end against a warm index (Supplementary S11).

A dedicated taxonomic-tree page at /tree renders any search result onto a linear cladogram of the underlying organism lineage, built from the resolved taxonomy table described under Data Sources and Integration, and visualizing seven taxonomic ranks from kingdom through compound. Queries returning more than one hundred compounds open with compound leaves hidden, and smaller result sets open with them visible. The default can be changed in the interface. Click-through on any clade node issues a refined search at the corresponding rank, and SVG and PDF exports preserve query terms and database version in the rendered image (Figure 2). The same query expanded to compound rank, and a broader region query rendered radially, appear in Supplementary Figures S13 and S14. Where a query cuts across taxonomy instead of within it, the same /tree route returns a multi-kingdom cladogram with separate subtrees coexisting in the result set.

**Figure 2.**
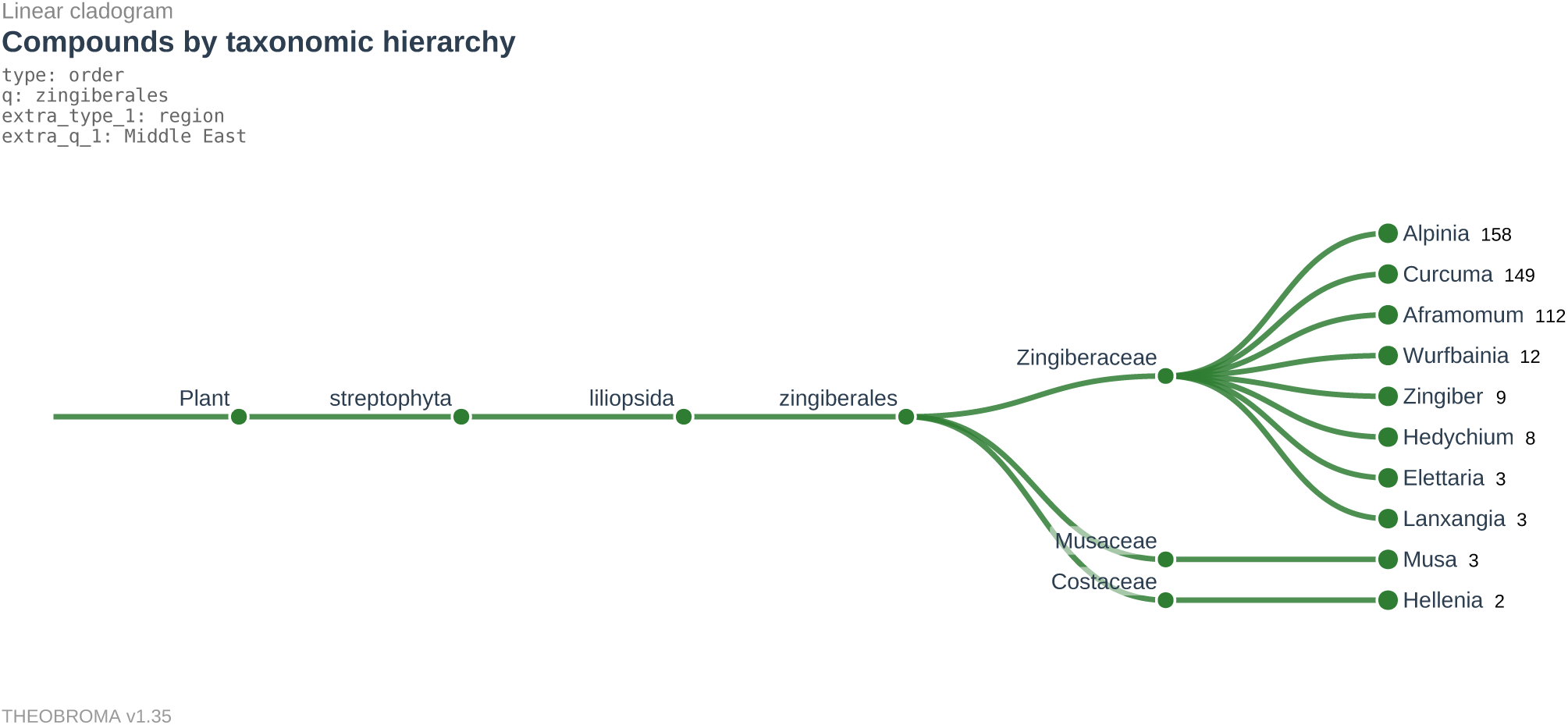
Linear cladogram returned by the THEOBROMA /tree route for a composed query combining taxonomic order (Zingiberales) and geographic region (Middle East). The 459 returned compounds distribute across three plant families (Zingiberaceae, Musaceae, Costaceae) and ten genera ranging from *Alpinia* at 158 compounds down to *Hellenia* at 2, illustrating the compositional power of the multi-filter /tree interface. Query terms, taxonomic hierarchy, and database version (v1.35) are baked into the PDF export.

Two persona-anchored scenarios illustrate the intended downstream workflow. An industrial medicinal chemist assembling a clean lead-generation library for commercial virtual screening filters at query time to the permissive-open plus public-domain tiers, returning the 900,103-compound subset through the /api/bulk endpoint with full annotation payloads. The alternative without per-compound license auditing would be manual re-audit of 29 source licenses for every commercial campaign. An academic natural-product researcher performing isomer-specific docking against the adenosine A_1_ and A_3_ receptors retrieves the *(Z,E)*-curcumin record (THEO 0213373) via the /api/stereoisomers/<COMP id> endpoint as a distinct family member rather than a collapsed duplicate. A binding differential between the two curcumin stereoisomers has been reported in this receptor context (29). The corpus holds the *(Z,E)* configuration only in a mono-anionic form, whereas the reported binding was measured on the neutral parents, so the retrieved record is the correct configuration in a different protonation state (Supplementary Figure S11). The alternative without stereochemistry-preserving deduplication would be hand-curation of literature-reported stereoisomers from primary sources.

Validation

Pre-deposit validation covered twelve categories from data integrity to adversarial input handling. Row counts match exactly between the CSV export and the database, with zero duplicate identifiers and zero null InChIKeys (Supplementary S11).

### Comparison with Peer Databases

While the peer cohort offers comparable scale, kingdom coverage, and modern stereochemistry-preserving deduplication, the open natural-product aggregator ecosystem otherwise converges on database-level license attribution, aggregate classification coverage without provenance stratification, and stereoisomer preservation without dedicated user-facing exposure. THEOBROMA addresses these three axes specifically. Table 1 summarizes THEOBROMA against the five most directly relevant open natural-product aggregators: COCONUT 2.0, LOTUS, the Natural Products Atlas 3.0, NPASS 3.0, and SuperNatural 3.0 (4; 5; 3; 6; 9). The comparison is restricted to the three differentiator axes.

Compound count is partly inherited, since four of the five peers named here are also integrated sources, but license resolution, classification provenance and stereoisomer exposure are properties of the integration and are not. The inherited fraction can be quantified. 1,012,047 compounds (89.3%) are attested by COCONUT or LOTUS and are in principle reachable through those two resources. The remaining 120,758 (10.7%) are not, contributed chiefly by FooDB (57,068), NPASS (14,194), HERB (11,849) and TM-MC (9,365) under primary attribution and not co-attested by either aggregator. A further 224,740 entries exist here as distinct records that COCONUT’s canonical-SMILES deduplication collapses, so the resolution gain is separable from the coverage gain.

Across the corpus 906,364 compounds (80.0%) carry a single attesting source and 226,441 carry two or more, and 137,149 of the latter (60.6%) span more than one license tier. Where COCONUT 2.0 and SuperNatural 3.0 aggregate annotation quality across axes into a single per-compound score, the classification tiers here stratify within one axis (4; 9), and where COCONUT 2.0 and IMPPAT 2.0 preserve stereochemistry internally through canonical isomeric SMILES or full-key designs (4; 8), the family relation is served through the /api/stereoisomers/<COMP id> endpoint and a radial display on every compound page. Two notes contextualize the table. COCONUT 2.0’s 695,133 unique compounds are counted under SMILES-based deduplication and are not directly comparable to THEOBROMA’s InChIKey-unique 1,132,805. LOTUS’s primary access channel is Wikidata with the legacy lotus.naturalproducts.net interface in a transitional retirement phase, and LOTUS counts vary by denominator across this manuscript (approximately 750,000 structure-organism pairs at the upstream Wikidata level, approximately 290,000 unique deduplicated structures at the peer-aggregator level, and 53,859 contributed into THEOBROMA after full-InChIKey deduplication and licensing audit in Supplementary Table S4, all defensible measures of different quantities). Table 1 is restricted to the three differentiator axes. Source counts, kingdom coverage, similarity search, geographic annotation, and ADMET content are compared in Supplementary Table S3.

### Discussion and Outlook

The per-compound license audit is the central design choice differentiating THEOBROMA from the peer cohort. License tracking at the resource level is established, with the Bioregistry recording per-resource licenses and the Reusable Data Project scoring license reusability (30; 31), but no open natural-product aggregator tracks licensing at compound granularity. Auditing at dataset granularity is established, notably the Data Provenance Initiative’s restrictive resolution across collections (32). The contribution here is resolution within a record, where 137,149 compounds carry attestations spanning more than one tier that no coarser scheme resolves. The consequence is operational. A relicensing event is bounded by the compounds its source attests, and a license-compatible subset can be reassembled by query rather than by re-audit. The two supporting axes operate at the access layer rather than the data layer. The peer cohort retains equivalent classification and stereochemical information internally, and the contribution here is its exposure as a queryable relation, which is what the abstract ranks below the licensing audit.

Three caveats remain. First, coverage of the provenance layers is uneven: geographic annotation reaches 23.3% of the corpus and taxonomic rank below kingdom 28.5%, both limited by whether the source organism matches a backbone entry in one of the three authorities. Kingdom coverage is higher, but only 327,906 compounds (28.9%) resolve against a backbone authority and the rest inherit their kingdom from source annotation, so the kingdom distribution is substantially inherited rather than independently resolved (Supplementary S2). Second, the classifier’s 26.5% model-inferred tier reflects Morgan-fingerprint discriminability limits on structurally atypical natural products and NPClassifier’s training coverage (33), and its reported macro-F1 is optimistic for unseen skeletons, since the train-test split is at compound rather than connectivity-family level (Supplementary S4). Third, the model tier over-predicts alkaloids. On the COCONUT-attributed subset the alkaloid share of the derived pathway rises from 25.4% among curated labels to 29.9% among NPClassifier assignments and 40.6% among model-inferred assignments. The gradient follows the provenance ordering, although the curated tier is not uniformly curator-assigned (Supplementary S4), and the 15-point margin between the curated and model tiers is consistent with the unweighted configuration selected under the macro-F1 objective (Supplementary Table S2). Class weighting was in the search space and was not selected. Recalibration against a class-balanced objective is deferred to a subsequent release. The persisted per-compound top-class probability and the provenance tier meanwhile let analysts impose their own acceptance threshold and exclude model-inferred assignments.

The v1.x roadmap covers four directions. Prediction extensions include protein-ligand interaction predictions from a forthcoming companion paper and formal retrospective validation of target-prioritization scores against held-out ChEMBL actives, reporting AUC-ROC and enrichment factors consistent with current target-prediction reporting standards (34). Source refresh will integrate ongoing updates from NPASS, LOTUS, and emerging regional resources such as GRAYU, BATMAN-TCM 2.0, and AlkaPlorer, with license re-audit at each integration cycle. Infrastructure work includes a community submission channel with versioned provenance tracking, expanded learned-embedding similarity search, and a versioned “commercial-safe” frozen export restricted to the CC BY 4.0 plus CC0 tiers (the 900,103-compound open-licensed subset) with its own DOI and frozen per-compound attestation snapshot, addressing the practical need for industrial downstream users to consume a static commercial-compatible release without re-running the per-compound license filter at query time. Multi-channel per-source licensing is also planned, so that both bounds account for sources distributing equivalent content under more than one set of terms. UI work includes explicit separation of taxonomic and chemical classifications and a navigable hierarchical view of the full ClassyFire and NPClassifier ontologies, both responding to user feedback on prior interface releases.

### Data Availability

The THEOBROMA v1.35 dataset is deposited on Zenodo under concept DOI https://doi.org/10.5281/zenodo.20443051, which resolves to the latest version, with this release at https://doi.org/10.5281/zenodo.21947744, released with per-compound source licenses preserved in the license tier field. The deposit is labelled v1.35.1, a packaging revision of the v1.35 data release described throughout. The corpus contents are identical. The CC BY 4.0 grant covers the contributed layer, namely the license audit and resolved tiers, the three-tier classification, the taxonomy resolution, the learned embeddings and the schema. Upstream terms flow through unchanged at compound granularity, so the aggregate is not uniformly CC BY 4.0 and downstream use should be governed by license tier rather than by the corpus-level statement. This article is released under CC BY 4.0 to maintain consistency with the licensing thesis of the work and the corpus-level license of the dataset. The reproducibility manifest (sources.yaml) is included in the Zenodo deposit and documents per-source download date, version, license, citation, and integration script for each of the 29 integrated sources. The live database is accessible at https://theobroma.l3s.uni-hannover.de with a REST API and OpenAPI specification documented at the same domain. L3S Research Center at Leibniz University Hannover has committed to a minimum five-year hosting period post-publication. Source code is maintained at https://github.com/ThorKlm/theobroma, and model artifacts plus derived datasets are mirrored at https://huggingface.co/ datasets/ThorKl/theobroma. Bulk downloads through the Zenodo deposit and the /api/bulk endpoint are available in CSV, JSON, and SDF formats. Per-query result sets are downloadable from the web interface in the same three formats plus tabular Excel.

## Acknowledgments

We thank the maintainers of the 29 source databases integrated into THEOBROMA for their continuing contributions to the open natural-products data ecosystem. We thank Chun-Wei Tung and colleagues for providing the TIPdb chemical structure archive. We thank the L3S Research Center at Leibniz University Hannover for production hosting and infrastructure support.

## Author Contributions

T.K.: Conceptualization, Methodology, Software, Validation, Formal analysis, Data curation, Investigation, Writing – original draft, Visualization, Project administration. A.J.: Methodology (taxonomic schema and phylogenetic-tree conceptualization), Validation (botanical and natural-product chemistry sanity checks), Writing – review and editing. J.F.: Methodology (natural-product chemistry guidance, classification and stereochemistry framing), Validation (chemical-class sanity checks, curcumin stereoisomer attribution), Writing – review and editing. W.N.: Supervision, Resources (L3S infrastructure and hosting), Writing – review and editing.

## Supplementary Material

This supplement provides methodological detail, per-category results, and supporting figures referenced from the main text, with figures placed at their point of reference. Section numbering mirrors the main-text flow: data integration and taxonomy (S1–S2), licensing audit (S3), annotation layers (S4–S6, S9–S10), web-interface access patterns (S7 compound result page, S12 batch annotation, S14 tree visualization), stereoisomer-family detail (S8), validation and infrastructure (S11), and the full comparison and per-source audit tables (S13).

## S1 Data Sources and Integration Detail

Each of the 29 sources was converted to a common schema through a per-source ingestion script recorded in the reproducibility manifest (sources.yaml), which documents download date, version, license, citation, source URL, and the integration script applied. Standardization with RDKit produced canonical isomeric SMILES, the full 27-character InChIKey, molecular formula, exact mass, and the standard Lipinski and medicinal-chemistry descriptor set. SMILES parsing succeeded on all entries of a stratified 10,770-compound validation sample (100%). 109 entries with NULL InChIKeys were recovered from the accompanying InChI string before re-validation. Salts are preserved as multi-fragment SMILES (5,443 entries, 0.5%) and isotopologues as distinct entries (303 compounds, 0.03%), reflecting a source-fidelity principle applied uniformly across annotation layers rather than an imposed normalization. Deduplication operates on the full 27-character InChIKey, which encodes stereochemistry and protonation state alongside molecular constitution. The 14-character connectivity prefix is retained as a grouping key for the stereoisomer-family relation. Standard InChI applies mobile-hydrogen normalization, so tautomer pairs differing only in the position of a mobile hydrogen share a connectivity prefix and fall in the same family, while tautomerism outside that layer is not normalized and yields distinct entries. The canonical isomeric SMILES generated alongside the key performs no tautomer normalization at all. Connectivity-prefix collisions between unrelated structures are documented but rare, and no attempt is made to detect them.

## S2 Taxonomy Resolution Methodology

Compound provenance is resolved through three authorities: the World Checklist of Vascular Plants (WCVP) for plant lineages, the NCBI Taxonomy database (13) for microbial and animal lineages, and the World Register of Marine Species (WoRMS) (14) for marine-organism cross-validation. Primary kingdom is assigned through a three-tier rebuild over the compound taxonomy attestations: an NCBI-plus-WoRMS majority vote where both authorities attest, WCVP-only resolution for plant attestations without microbial or animal evidence, and a source-level fallback where authority evidence is absent. A 510-family backfill following APG IV (35) and PPG I (36) plant-lineage circumscriptions, extended to gymnosperm, bryophyte and green-algal families outside both, recovers kingdom assignment for compounds whose source organism resolves to family rank but not to a sequenced taxon, and a 75%-threshold consistency pass reassigns compounds whose attestations disagree below the majority threshold. The backfill table covers 448 angiosperm families, following APG IV circumscriptions where they apply, 29 fern families following PPG I, and 33 further gymnosperm, bryophyte and green-algal families outside both circumscriptions. The three tiers contribute unequally. 355,970 compounds (31.4%) carry at least one organism attestation in compound taxonomy, of which 327,906 (28.9%) resolve against a backbone authority to an accepted taxon. The 804,899 compounds (71.1%) without a backbone-resolved taxon receive their kingdom through the source-level fallback, 736,704 of them attributed to COCONUT. Kingdom coverage across the corpus therefore rests substantially on source-provided annotation, not on independent lineage resolution, and the genus and family ranks are populated only for the resolved fraction: 323,402 and 323,091 compounds respectively.

Primary kingdom is assigned by majority vote across attestations and reconciled against the kingdom implied by the resolved phylum, so that no compound carries a kingdom contradicting its own lineage. Kingdom filtering matches the resolved kingdom or any secondary kingdom recorded in secondary kingdoms, so a compound reported once from a fungus is returned under a fungal query even where its resolved lineage is plant: of the fungal hits, 41,545 match on the primary assignment and 7,367 on a secondary attestation alone. The recorded secondaries are a subset of the kingdoms present in the underlying attestation chain, which the compound page exposes in full. Compounds attested across three or four kingdoms (5,152) typically carry twenty or more source organisms and reflect the breadth of upstream literature aggregation rather than confirmed cross-kingdom occurrence. Secondary attestations are marked in kingdom search results, and the full per-source attestation chain is shown on the compound page.

The resolution retires the v1.0 “marine” pseudo-kingdom: previously marine-tagged compounds are redistributed by actual lineage to animal (predominantly sponges, corals, and mollusks), fungi (marine ascomycetes), plant (marine algae), bacteria (cyanobacteria), and unresolved. Compounds in protist supergroups (Stramenopiles, Alveolata, Rhizaria, Hacrobia, and Excavata, spanning the phyla Oomycota, Bacillariophyta, Ochrophyta, Ciliophora, Myzozoa, Haptophyta, Euglenozoa, Foraminifera, Discosea, Cercozoa, and Bigyra) are assigned to unresolved rather than forced into a canonical kingdom, since they are biologically protists rather than plant, animal, fungi, or bacteria. The resulting kingdom distribution and the secondary-kingdom cross-attestation breakdown are given in Table S1.

The 486,032 stereoisomer families span a long-tailed size distribution: 171,290 are singletons with no stereochemical or protonation variation, 198,112 contain two members, and 6,632 hold ten or more members. The curcumin connectivity skeleton is one example, resolving to eight distinct full-key entries: two parent forms in the neutral protonation state (InChIKey suffix ‘-N‘, one encoding the (*E*,*E*)- configuration and one carrying no stereochemical layer), and six protonation-state variants in the mono-anion (‘-M‘) and di-anion (‘-L‘) states spanning three stereochemical configurations. The third configuration is observed only in the anionic forms. This asymmetry between neutral and protonated stereochemical coverage is an artifact of InChI stereochemical perception observed here, which can assign different stereodescriptor coverage to neutral versus deprotonated forms of the same connectivity skeleton. Most connectivity skeletons appear in one or two stereoisomeric forms, with a small minority showing extensive variation through combined stereochemical and protonation state.

## S3 Licensing Audit Methodology

License metadata was extracted per source in order of precedence: the source’s Zenodo or institutional repository, the peer-reviewed citation, the public download or terms-of-service page, and direct correspondence with maintainers where the preceding sources were silent or ambiguous. Each source was assigned to one of five tiers: permissive open (CC BY 4.0 or equivalent), public domain (CC0), non-commercial (CC BY-NC 4.0 or equivalent), copyleft (share-alike licenses such as CC BY-SA or CC BY-NC-SA), and unspecified (no explicit data-level license). Where the data-level license was ambiguous rather than absent, the most restrictive defensible upstream signal was adopted. Four sources (NPASS, phytochemdb, CSIRO, and ANPDB) were assigned to the non-commercial tier on this basis, since their data-level terms were not stated explicitly and the strongest available signal (a non-commercial paper license, a commercial institutional origin, or a regional repository without an explicit data license) did not support a permissive-open designation.

Two sources distribute through more than one channel and merit explicit treatment. COCONUT 2.0 carries CC BY 4.0 on the Zenodo deposit from which THEOBROMA ingested it (record 13692394, downloaded September 2024) and CC0 on the project download page, which serves a rolling current release rather than the September 2024 snapshot used here. LOTUS carries CC BY 4.0 on the Zenodo export ingested here and CC0 on Wikidata, which is a separate retrieval channel rather than the same artifact under different terms. In both cases the license map records the designation attaching to the artifact actually retrieved. COCONUT’s terms of service further state that it places no restrictions on redistribution beyond those provided by the original data owners, so its CC0 designation covers COCONUT’s own contribution rather than clearing the terms of the sources it aggregates. Share-alike sources were excluded from integration to avoid relicensing the aggregate: CMNPD (CC BY-NC-SA 4.0) was excluded under the conservative operational interpretation adopted here, since ShareAlike would require the aggregate to be released under compatible terms if the integration were treated as an Adaptation rather than a Collection. The NonCommercial term alone keeps CMNPD outside the commercially reusable fraction independently of the ShareAlike question. Future inclusion of share-alike sources would require a separately licensed share-alike aggregate. Creative Commons itself discourages NonCommercial and NoDerivatives terms on scientific data and recommends CC0 for maximal reuse, which is consistent with the finding here that non-commercial attestation is the dominant constraint on the aggregate.

The per-source assignment described above is the first of two stages. The second stage resolves each compound’s license tier across all attesting sources under a most-restrictive-wins rule. The rule is a conservative operational choice rather than a legal determination: where a compound is attested by sources whose terms differ, honoring the most restrictive attestation is the interpretation under which no source’s terms can be breached, and the combined obligation is the union of the individual ones. The audit records the terms each maintainer asserts over the data it distributes. It does not adjudicate whether those terms are enforceable over an individual record, which depends on jurisdiction and on whether copyright, sui generis database right, or contract is the operative instrument. Users with specific redistribution requirements should treat the resolved tier as a compliance floor rather than as legal advice.

Precedence is ordered, from most restrictive to least restrictive, as: copyleft, unspecified, non-commercial (CC BY-NC 4.0), permissive open (CC BY 4.0), and public domain (CC0). Copyleft is placed at the most-restrictive position because share-alike obligations propagate to derivatives. Unspecified is treated as the second-most restrictive because the absence of a documented license should not grant more permissive use than the most conservative defensible interpretation. The precedence rank and the practical redistribution floor answer different questions and should not be conflated. The rank governs combination: an unspecified attestation is not overridden by a permissive co-attestation. The floor governs use: no defensible interpretation of an undocumented license grants more than non-commercial, so a downstream user should treat unspecified as non-commercial-equivalent. Both point in the same conservative direction, and the tier is kept separate rather than merged into non-commercial so that the absence of documentation remains visible in the audit. Non-commercial sits above permissive open because the non-commercial-only restriction is a stronger constraint than attribution alone. Permissive open sits above public domain because a CC BY 4.0 attestation imposes an attribution requirement that CC0 does not, and a compound co-attested by both must carry attribution to comply with the CC BY 4.0 source. A compound attested only by CC0 sources resolves to CC0 because no attribution-requiring source claims it. The same precedence ordering applied in the opposite direction gives the least restrictive attested tier among the artifacts ingested, one license per source. This is not a redistribution license for the aggregate: it records that the compound is available under those terms from at least one integrated source, and a user relying on it should obtain the compound from that source and attribute it rather than THEOBROMA. Where a source distributes equivalent content through a further channel, the terms of that channel lie outside the audit and remain available directly. LOTUS is the case in the present release, publishing on Wikidata under CC0 alongside the CC BY 4.0 Zenodo export recorded here, which affects the 226,699 compounds it attests. Recording multiple distribution channels per source is planned for a subsequent release.

Resolution proceeds as follows for each compound in the corpus:

1. Parse the compound’s all sources string into the set of attesting sources.
2. For each attesting source, look up the per-source license from the audit-derived mapping (29 sources, persisted as license map.tsv alongside the reproducibility manifest).
3. Apply the precedence ordering and return the most restrictive (lowest-rank) tier across the attestation set.
4. Write the resolved tier to compounds.license tier, persist the attestation chain in per source license attestation for downstream audit.

The attestation chain is materialized at ingestion as a dedicated per source license attestation table with one row per (compound, attesting source) pair, recording the source’s license at the moment of ingestion and the timestamp of attestation. The materialization decouples the per-source license vocabulary from query-time evaluation and provides a structured audit trail that survives subsequent edits to the source-license mapping. In v1.35, the table contains 1,512,594 rows across 1,132,805 unique compounds, giving 1.34 attestations per compound corpus-wide. The 431,099 compounds carrying a non-null primary name are a distinct denominator from the 149,178 synonym- bearing compounds that give the synonyms-per-compound figure of 3.23 reported in the main text. Per-source contributions under primary attribution span four orders of magnitude and are shown by license tier in Figure S2.

The resolver is implemented as a single idempotent SQL transaction (scripts/apply license resolver.sql in the repository) and produces the same output on any re-run against the same attestation table. Applied to v1.35, the resolver reassigns 103,304 compounds (9.1% of the corpus) from the per-source first-stage assignment to a more restrictive tier on the basis of multi-source co-attestation. The dominant transition is from permissive open to non-commercial (94,940 compounds co-attested by a CC BY-NC 4.0 source alongside a CC BY 4.0 source), followed by smaller transitions into the unspecified tier (4,657 compounds across three transitions involving an unspecified-license co-attestation: 2,523 from permissive open, 2,043 from non-commercial, and 91 from public domain), the public-domain-to-non-commercial transition (2,818 compounds where TM-MC attestations co-occur with a non-commercial source), and the internal redistribution from public domain to permissive open under the attribution-requirement rule (889 compounds where TM-MC attestations co-occur with a CC BY 4.0 source). The headline post-resolver tier distribution reported throughout the manuscript (891,860 / 225,536 / 8,243 / 7,166 across permissive open, non-commercial, public domain, and unspecified respectively, with 900,103 compounds resolving to open-licensed status) is a direct database reading of the compounds.license tier column. Applying the transitions above to the per-source tier totals of Supplementary Table S4 reproduces it exactly. The 988,434 compounds whose primary source is permissive open lose 97,463 to more restrictive tiers and gain 889 from public domain, giving 891,860. Public domain, non-commercial and unspecified close in the same way.

The resolved per-compound tier is exposed alongside the underlying attestation chain through the /api/compound/ <ID>/license-provenance endpoint, returning the resolved tier, the resolution rule, the precedence ordering, and the per-source attestation list with each source’s license. Downstream users can audit any compound’s resolution independently of the manuscript or the database documentation, and the same endpoint supports bulk compliance workflows by composing with the corpus-level license filter of the main interface.

## S4 Classification Methodology

The classifier is documented here as annotation infrastructure rather than as a methodological contribution. The hyperparameters and evaluation are reported so that the assignment of a class to any compound in the model tier can be reproduced, not as a claim about classifier performance in general. Class assignments carried in the source databases are accepted directly for the curated tier, and NPClassifier assignments for the tool tier. The tier records where an assignment came from rather than how it was produced. Several sources, COCONUT among them, generate their own NPClassifier and ClassyFire annotations, so a source-supplied class is not necessarily curator-assigned. Superclass and pathway are derived from class through the NPClassifier ontology and materialized in effective superclass and effective pathway. The derived pathway is the column reported in Figure S3 and throughout this manuscript. The source-supplied np pathway column is retained unmodified but is not used, because it disagrees with the class of the same record for 43,656 compounds on the curated tier by assigning Alkaloids where the class resolves elsewhere, against 2,041 disagreements in the opposite direction. The defect predates v1.34 and originates in the initial ingestion. Compounds whose np pathway was written by the NPClassifier run of this project rather than inherited show no such divergence. The two columns also differ in coverage. The derived pathway reaches 1,132,653 compounds against 1,101,638 for the source column, since 28,640 model-tier compounds carry a derived pathway but no source-supplied one and 2,512 source-inherited classes arrived without an accompanying pathway, while 6 unclassified compounds carry a source pathway but no class and 131 carry a source pathway whose assigned class does not resolve to a derived one. The 152 compounds without a derived pathway comprise the 21 unclassified and 131 whose assigned class does not resolve to a pathway under the primary-parent rule. The remaining compounds are classified by a per-class XGBoost ensemble operating on a 2,471-dimensional feature vector that concatenates Morgan fingerprints (37), MACCS keys, and PCA-projected ChemBERTa embeddings. Compounds whose curated label spans multiple classes are excluded under the single-label formulation, labels traceable to the artifact-suspect upstream set are excluded, and classes with fewer than 50 training examples are dropped to avoid unstable low-support predictions. The resulting labelled pool comprises 384,006 distinct compounds across 501 of NPClassifier’s 672 classes. A stratified 15% holdout of 57,601 compounds is reserved first, and the remainder is split again into 267,652 training and 58,753 validation compounds, with five-fold cross-validation performed within the training set. The split is at compound level, not connectivity-family level: 80% of labelled compounds share a 14-character prefix with another, and 78% of those families carry a single class, so the reported macro-F1 is optimistic for unseen skeletons. Grouped-split evaluation is deferred to the next update. Hyperparameters were selected via Optuna against a 5-fold cross-validated macro-F1 objective, with the search space and selected values reported in Table S2. Classification quality is summarized by macro-F1 (averaged across classes, sensitive to rare-class performance) and micro-F1 (aggregated across instances, dominated by common classes), both reported with the per-class confusion structure in the project repository. No acceptance threshold is applied at assignment time. The per-compound top-class probability is persisted so that a threshold can be imposed downstream, and the twenty-one compounds left unclassified are those for which no feature vector could be computed rather than those falling below a confidence floor.

## S5 Chemical Descriptors and Embeddings Detail

Learned embeddings are 768-dimensional mean-pooled representations from the original ChemBERTa checkpoint seyonec/ChemBERTa-zinc-base-v1 (Chithrananda et al. 2020, arXiv:2010.09885, RoBERTa-base architecture pre-trained on the ZINC-250k SMILES corpus, uploaded to HuggingFace April 2020), computed per compound and indexed via a FAISS HNSW graph for sub-linear nearest-neighbor retrieval over the full corpus. The two later models in the ChemBERTa family (ChemBERTa-2, Ahmad et al. 2022, arXiv:2209.01712, and ChemBERTa-3, Singh et al. 2026, Digital Discovery 5, 662, DOI 10.1039/D5DD00348B) are independent training frameworks with different architectures and pretraining data and are not used in THEOBROMA v1.35. A second learned representation is provided by NaFM (28), a graph isomorphism network pretrained on natural products through scaffold-subgraph reconstruction and scaffold-aware contrastive learning, in contrast to the general-chemistry pretraining corpus underlying ChemBERTa. Embeddings are 1,024-dimensional and indexed in a half-precision flat FAISS index, and the ChemBERTa index uses HNSW with *M* = 32, efConstruction = 200, and efSearch = 128. NaFM retrieval is available for structures resolvable to a corpus entry by identifier, name, or InChIKey and is not computed at query time for off-corpus SMILES. NaFM’s node features encode chirality alongside atom type and formal charge, so the two learned representations differ in stereochemical sensitivity: the ChemBERTa tokenizer does not distinguish chiral notation, while NaFM does. Both embeddings serve nearest-neighbor retrieval only. The classifier feature vector of Supplementary S4 uses the ChemBERTa projection alone. Embedding geometry separates established structural classes cleanly, with a silhouette coefficient of 0.633 measured across aromatics, xanthines, salicylates, and steroids. Physicochemical descriptors (exact mass, logP, topological polar surface area, hydrogen-bond donors and acceptors, ring count, rotatable bonds) are computed with RDKit. Three structural-alert sets are applied as published: PAINS for pan-assay interference, BRENK for unstable or reactive moieties, and the NIH filter set. Bemis-Murcko scaffolds and Ertl-Schuffenhauer synthetic accessibility scores complete the per-compound descriptor surface. The synthetic accessibility distribution across the corpus is shown in Figure S4.

## S6 Traditional-Medicine Tagging Detail

Traditional-medicine provenance is propagated from source-database attribution, with HERB 2.0, TM-MC 2.0, and IMPPAT 2.0 as the primary contributors and additional tags accruing through shared compound identity across sources. The raw tagging vocabulary reflects source-specific labeling conventions rather than a single imposed ontology, spanning Traditional Chinese Medicine and related East Asian systems and Ayurveda and Indian traditional medicine, with composite labels (for example, “TCM / Korean / Japanese traditional medicine”) preserved as attested. A compound attested by sources belonging to more than one tradition after cross-source propagation is counted as multi-tradition. 4,841 compounds carry such cross-cultural attestation (8.0% of the 60,242 tagged compounds), documenting shared use of widely distributed medicinal plants. Note that the figure-level “tag count” metric used in Figures S5, S6, and S7 is a slightly broader set than “multi-tradition”: because composite source labels (for example “TCM / Korean / Japanese traditional medicine”) count as multiple tags before propagation, the subsample tag-count-at-least-two fraction reaches approximately 11% within tagged compounds, above the corpus-wide multi-tradition rate of 8.0%. THEOBROMA inherits the access and benefit-sharing status of its upstream sources in the spirit of the Nagoya Protocol. Commercial development on traditional-medicine-tagged compounds should additionally verify Nagoya-relevant compliance against the originating source.

The same embedding admits a complementary validation view. Figure S6 colors a uniform 50,000-compound subsample by effective pathway, showing that the mean-pooled ChemBERTa representation organizes natural-product chemical space along biosynthetic lines: the two largest categories occupy coherent regions and the smaller ones resolve into their own zones rather than dispersing uniformly, which is consistent with the use of this embedding for the nearest-neighbor similarity search described in the main text. The subsample is uniform rather than tag-stratified, so pathway proportions track corpus prevalence. The sparsest pathway, carbohydrates at 533 compounds in the subsample, is correspondingly underrepresented.

A further view of the same uniform subsample by source-organism kingdom (Figure S7) shows that the embedding also resolves taxonomic structure. Plant-derived compounds form the majority of the subsample (85.2%) and span the embedding broadly, consistent with their wide chemical diversity. Smaller kingdoms (animal 7.1%, fungi 3.6%, bacteria 2.8%) and unresolved-origin compounds (1.3%) distribute across the same chemical space, with kingdom-specific concentrations visible where biosynthetic repertoires diverge from the plant baseline (for example bacterial polyketide-rich regions, fungal alkaloid clusters).

## S7 Compound Result Page

The per-compound result page at /compound/<COMP id> is the canonical single-compound view, rendering the full annotation surface into a single scrollable layout that culminates in an interactive radial display of the stereoisomer family. Figures S8, S9, and S10 show the rendered page for fisetin (THEO 0858442) split across three captures, since the integrated layout exceeds two A4 pages at readable scale. The same layout applies to every compound in the corpus. Per-compound variation reflects which annotation layers carry populated values for a given structure (traditional-medicine tags appear only where the compound is attested in a pharmacopoeial source, ADMET predictions appear for all 1,132,789 RDKit-processable compounds, and the stereoisomer-family radial renders when the connectivity prefix has at least two attested members).

## S8 Stereoisomer Family Detail

The 486,032 stereoisomer families span a long-tailed size distribution: 171,290 are singletons with no stereochemical or protonation variation, 198,112 contain two members, and 6,632 hold ten or more (for example, the curcumin connectivity skeleton resolves to eight distinct full-key entries). The deduplication at the full 27-character InChIKey preserves binding-relevant stereochemical and protonation distinctions that 14-character truncation would erase. Figure S11 illustrates this for the (*E*,*E*)- versus (*Z*,*E*)-curcumin pair.

## S9 Target Prioritization Detail

The per-compound score vector is intended as a structure-based browsing affordance for candidate triage and library exploration, exposed in the API and compound-detail page as an exploratory feature. The scores carry no calibrated probability interpretation, the panel selection itself reflects the exploratory interests of the THEOBROMA team rather than systematic coverage of validated assay targets, and the absence of decoy-set evaluation means the scores cannot be interpreted as enrichment estimates. A qualitative consistency check of the highest-scoring compound per panel recovers structurally plausible representatives across all 25 panels (Erlotinib for EGFR, Mozenavir for HIV-1 protease, and Paclitaxel for *β*-tubulin among the named exemplars), confirming that the maximum-Tanimoto operation behaves as expected without making any quantitative claim about predictive performance.

## S10 Trust Score Definition

The trust score quantifies per-compound annotation coverage across nine axes: structure, classification, scaffold, taxonomy, geography, traditional-medicine tagging, ADMET prediction coverage, novelty assessment (defined as the Tanimoto distance from the compound to the nearest FDA-approved drug in the DrugBank-curated approved set (39), normalized to [0,1] with higher values indicating greater structural distance from established drug-like chemistry), and similarity-to-bioactive scoring. Each axis contributes a normalized component on a 0–1 scale under a configurable weight vector, and the default-weight corpus mean is 0.326, corresponding to roughly three of the nine axes annotated per compound on average. This low mean is a structural property of open natural-product aggregation rather than a quality deficiency: source databases typically contribute to a few axes each (a chemistry-curated source provides structure and classification, an ethnobotanical source provides taxonomy and traditional-medicine tags, an ADMET-prediction layer covers its endpoints alone). Per-compound coverage therefore saturates only at the union of all contributing sources, which is uncommon for any individual compound.

## S11 Validation and Latency Benchmarks

Pre-deposit validation was performed across twelve categories: data integrity, cross-table consistency, license-tier integrity, reference-compound lookup, traditional-medicine tagging, novelty reproducibility, target-prediction sanity, API and web-interface behavior, external link resolution, README accuracy, tester feedback, and adversarial input handling. Row counts match exactly between the CSV export and the database at 1,132,805 compounds, with zero duplicate identifiers and zero null InChIKeys. A stratified 10,770-SMILES sample parsed at 100% and nineteen adversarial test cases across six input-handling categories (malformed SMILES, oversized queries, injection-pattern strings, empty and whitespace inputs, out-of-range parameters, and unsupported content types) all returned correctly coded client errors or empty result sets without server-side faults.

Query latency was measured on the production node (8 vCPU AMD EPYC 7662, 32 GB RAM, no GPU, warm PostgreSQL cache and warm similarity indices). Reported values are end-to-end served response times, including HTTP handling and the serial request queue, rather than isolated index-search cost. Median served latency is 40 ms for textual API search, 590 ms for Morgan and 700 ms for MACCS fingerprint Tanimoto similarity, 630 ms for NaFM embedding similarity over the half-precision flat index, and 970 ms for ChemBERTa embedding similarity over the HNSW index. Each figure was obtained twice by independent measurement and the two agree. Medians rather than tail percentiles are reported because the service runs a single worker holding the similarity indices in process, which serializes concurrent requests. Substructure search is available through the web interface rather than a dedicated API endpoint and was not included in this measurement.

## S12 Batch Annotation Endpoint

The /annotate page and corresponding /api/annotate REST endpoint accept structure-list uploads in CSV, TSV, Excel, plain text, or SMILES (.smi) format and return per-compound annotation for corpus intersections. Column auto-detection identifies SMILES or InChIKey columns from either whitelisted header names (smiles, inchikey, and common synonyms) or value-pattern sniffing across the first twenty data rows. Files without a header row are accepted with synthetic column-name generation. The page chunks inputs at 500 per call to keep individual round trips bounded, exposes a determinate progress bar, and produces two downloadable CSVs (matched and unmatched). Each matched row returns the full annotation surface: compound identifier, source attribution, license tier, three-tier classification fields, ADMET context, taxonomic aggregation (phyla, classes, orders, families, genera), and the stereoisomer-family-member list under the 14-character InChIKey connectivity prefix. Unmatched rows carry the original identifier and SMILES with a per-row reason code. Processing is by lookup against the precomputed corpus rather than on-the-fly prediction on arbitrary uploads, keeping the endpoint within the scope of a data resource rather than introducing new computational results.

The batch upload-and-enrich pattern itself is established prior art. The EPA CompTox Chemicals Dashboard Batch Search is the canonical analogue, accepting identifier uploads at scale with column selection and multi-format export (40). Related single-call enrichment services include the PubChem Identifier Exchange Service, SwissADME, the Chemical Translation Service, and UniChem, each operating across their own chemistry domains. The specific contribution of THEOBROMA’s endpoint is therefore not the batch-enrichment pattern itself but its instantiation against a curated natural-products corpus with two NP-relevant additions: the license tier returned per matched compound, which composes with the corpus-level license filter to assemble license-compatible subsets at batch scale, and the stereoisomer-family-member list returned alongside each match, which surfaces the same 14-character connectivity-prefix family exposed via /api/stereoisomers/<COMP id> without requiring per-compound lookup.

Within the open natural-product subfield, no peer aggregator currently offers an equivalent upload-and-enrich endpoint. COCONUT 2.0 provides single-structure search, bulk corpus download, and a REST API but no upload-and-enrich page (4). Natural Products Atlas 3.0 exposes a REST API without a batch-upload interface (3). LOTUS provides bulk access through SPARQL on Wikidata, with the learning curve acknowledged by its authors (5). NPASS 3.0 accepts batch identifier queries but without file upload, column auto-detection, or unmatched-set reporting (6). The gap is in the NP-specific instantiation of an otherwise established access pattern. The contribution is filling it with the licensing- and stereoisomer-aware additions above. Figure S12 shows the workflow on a four-compound test input.

## S13 Full Comparison and Per-Source Audit Tables

This section carries the full versions of the two tables abridged in the main text. Table S3 extends the main-text peer comparison beyond the three differentiator axes to source count, kingdom coverage, similarity search, geographic annotation, and ADMET content. Table S4 gives the per-source licence audit across all 29 integrated sources, with the compound-level resolved-tier summary in the notes.

## S14 Taxonomic Tree Visualization at Scale

The /tree route described in the main text varies its display depth with result-set size and offers both linear and radial layouts. Two cases bracket its range. Figure S13 shows the maximum-depth case: the cladogram for a composed bacterial query (Aminoglycosides in the 400 to 600 Da band) expanded to the individual compound leaves beneath each genus, with compound names and identifiers rendered at the terminal nodes. This complements the genus-rank rendering in the main text (Figure 2), where the larger result set leaves compound leaves hidden by default. Figure S14 shows a broader single-axis query (region restricted to Africa) rendered radially, where the taxonomic hierarchy across the represented kingdoms and phyla remains legible because the radial layout distributes nodes around the circumference rather than competing for horizontal space at a single depth. Together the two figures span the route’s range, from per-compound resolution on a narrowly composed result set to a radial overview of a broad single-filter query, and between them span plant and bacterial lineages and all four resolved kingdoms.

---

**Figure S1.**
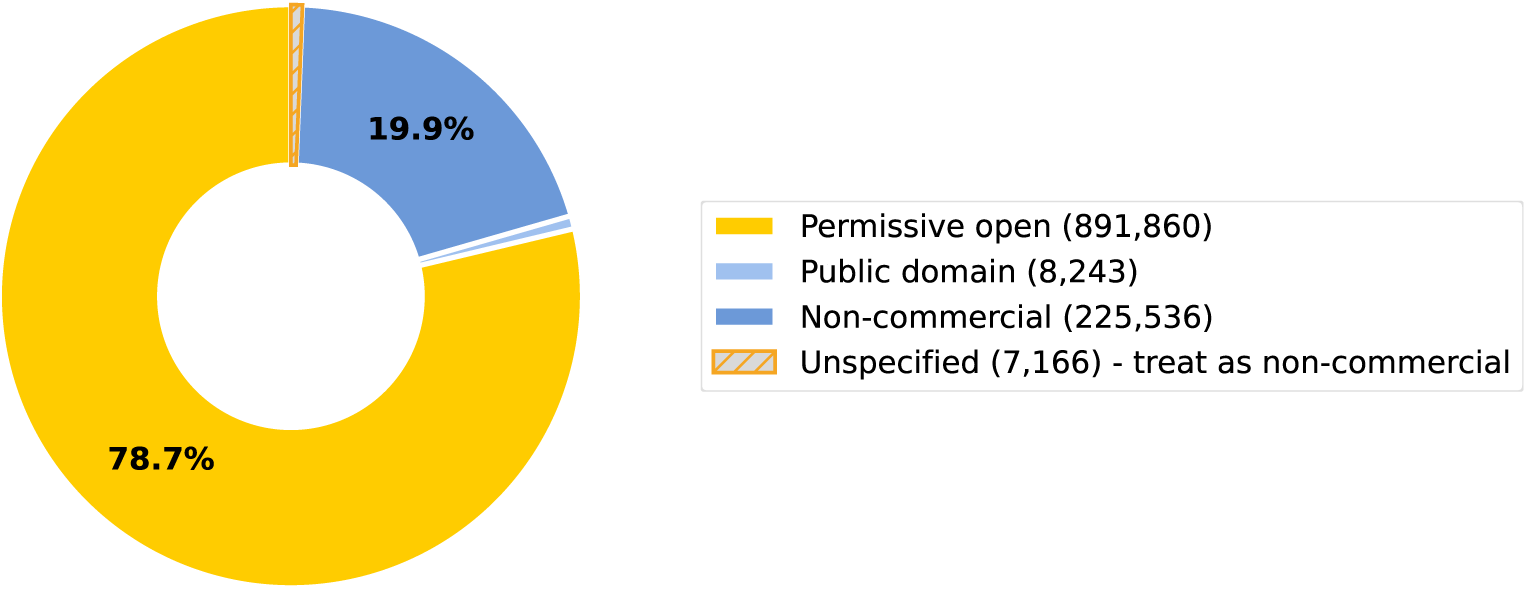
License-tier distribution across 1,132,805 compounds after the most-restrictive-wins resolver has been applied: permissive open 891,860 (78.7%), non-commercial 225,536 (19.9%), public domain 8,243 (0.7%), unspecified 7,166 (0.6%). The open-licensed fraction (permissive open plus public domain) is 900,103 compounds (79.5%). The unspecified wedge is rendered separately rather than merged into non-commercial, so that the audit signal survives at the visualization layer. Its precedence position is explained above. The least restrictive bound is reported separately in the Licensing Audit section.

**Figure S2.**
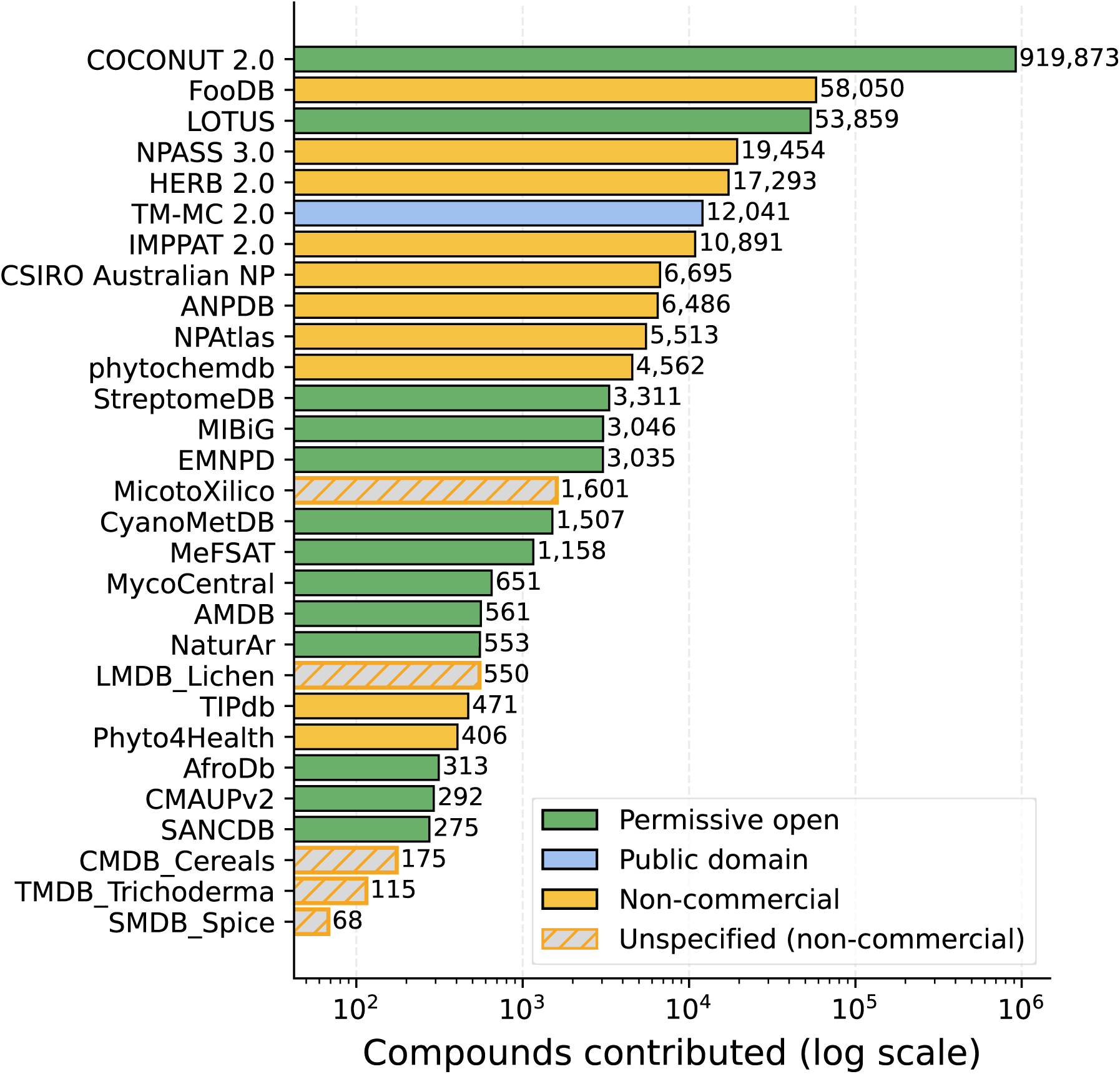
Rendered on a log scale and colored by license tier, per-source compound contributions span four orders of magnitude, from COCONUT 2.0 at 919,873 compounds under primary attribution to SMDB Spice at 68. Values are those of Table S4 and reflect primary attribution rather than total attestation.

**Figure S3.**
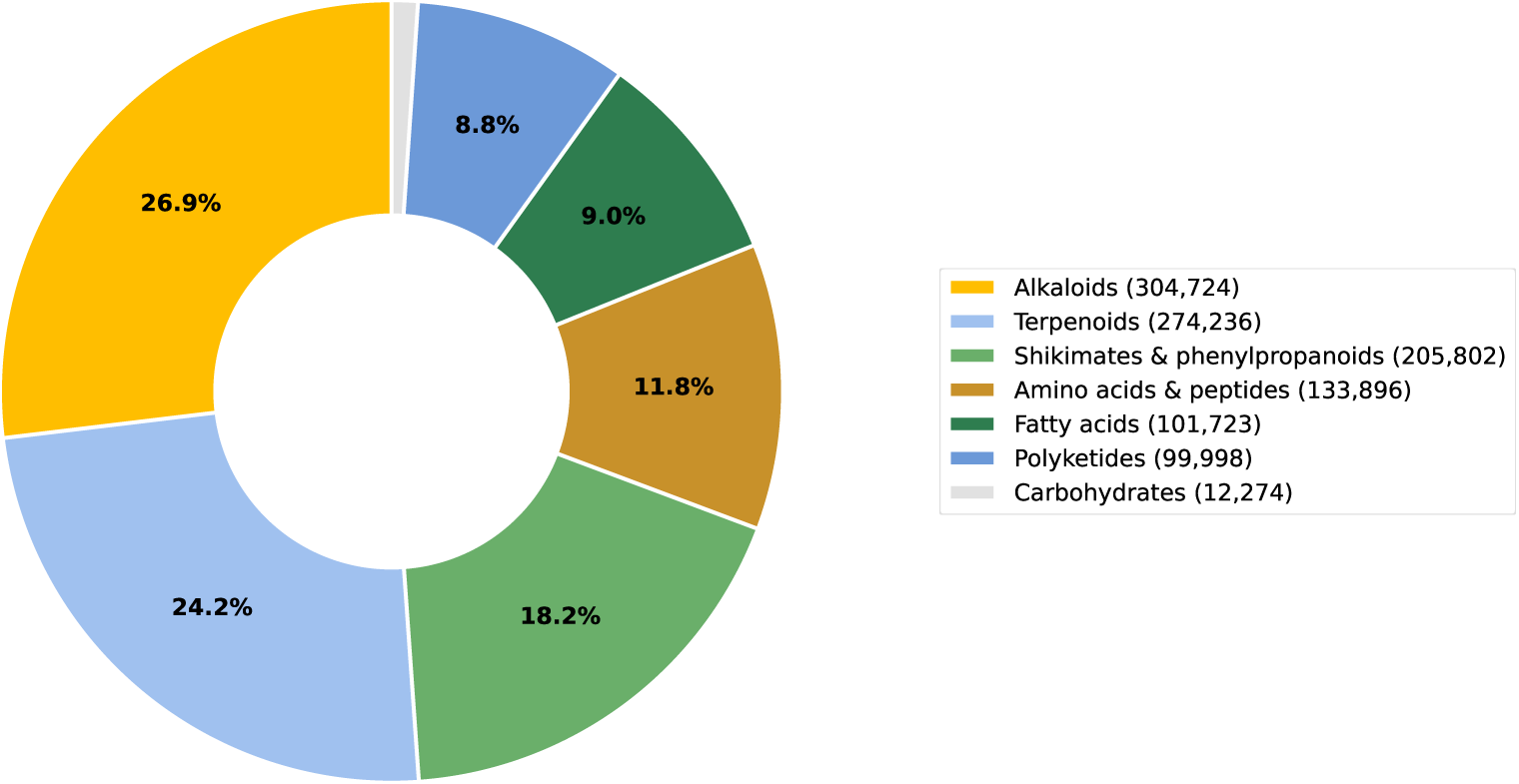
NPClassifier pathway distribution across the 1,132,653 compounds carrying a derived pathway, taken from effective pathway, which is resolved from the assigned class through the NPClassifier ontology under a designated primary parent. Wedge labels are shown for shares above 3% and all seven values are given here: alkaloids 304,724 (26.9%), terpenoids 274,236 (24.2%), shikimates and phenylpropanoids 205,802 (18.2%), amino acids and peptides 133,896 (11.8%), fatty acids 101,723 (9.0%), polyketides 99,998 (8.8%), carbohydrates 12,274 (1.1%). The source-supplied np pathway column is not used here. On the curated tier it disagrees with the class of the same record for 43,656 compounds by assigning Alkaloids where the class resolves elsewhere, against 2,041 disagreements in the opposite direction (Supplementary S4). The NPClassifier pathway distribution across the pathway-annotated corpus is shown in Figure S3.

**Figure S4.**
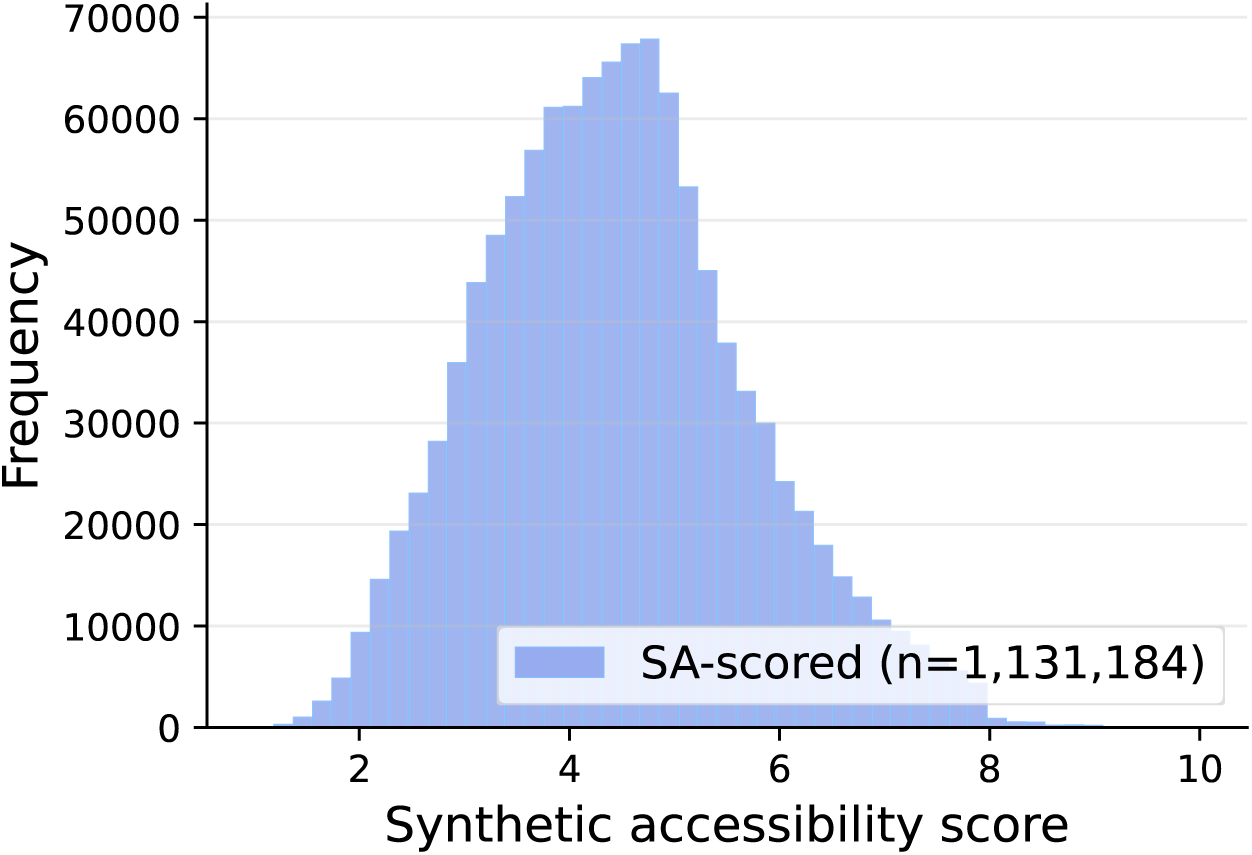
Synthetic accessibility score distribution (Ertl-Schuffenhauer) across the 1,131,184 compounds for which a score could be computed, 99.86% of the corpus.

**Figure S5.**
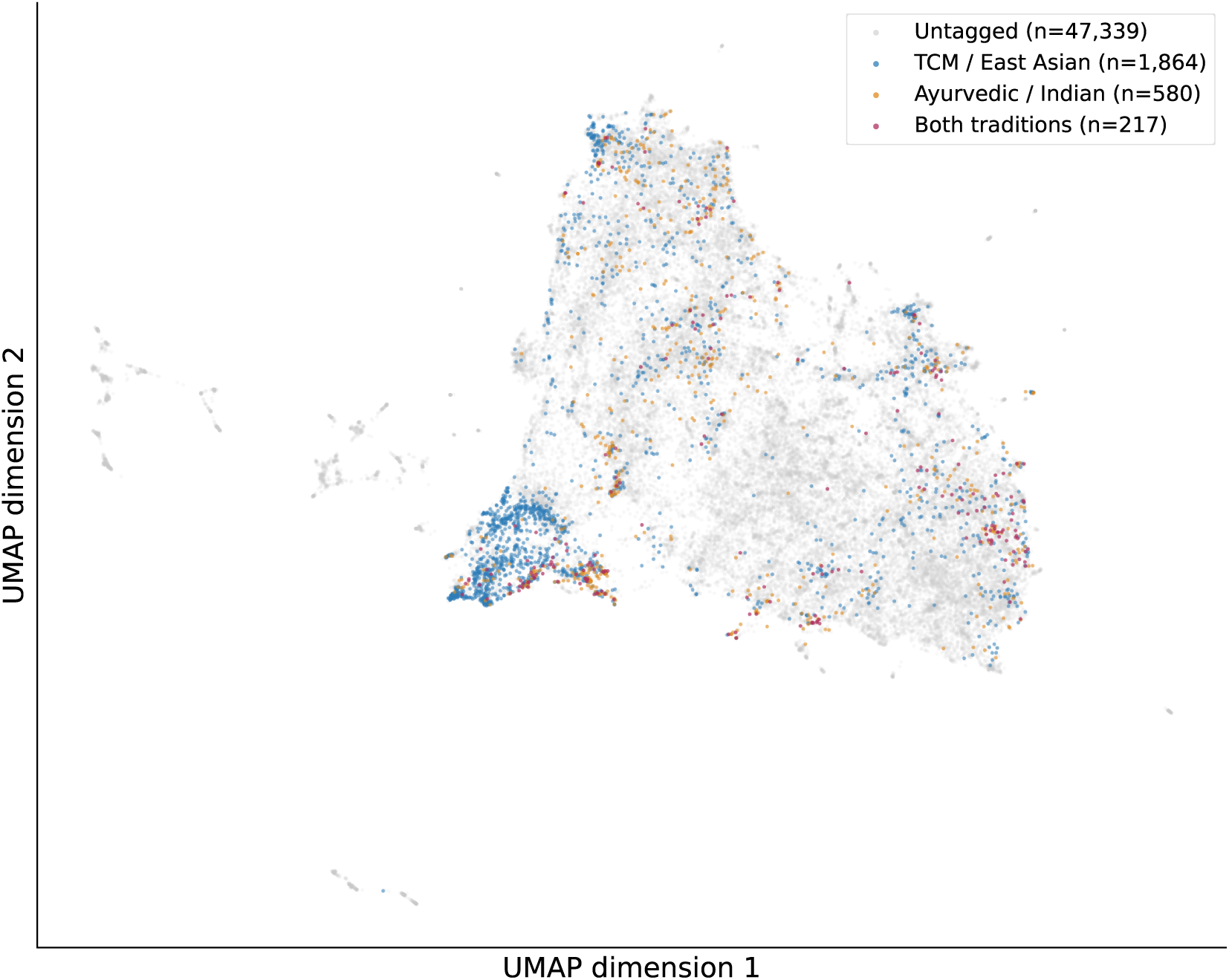
ChemBERTa embedding space colored by traditional-medicine attestation. UMAP projection of mean-pooled 768-dimensional ChemBERTa embeddings (seyonec/ChemBERTa-zinc-base-v1, n_neighbors_ = 15, min dist = 0.1, cosine metric, fixed random state) across a uniform 50,000-compound subsample. Compounds are colored by the tradition under which they are attested after cross-source propagation: untagged (47,339), Traditional Chinese Medicine and related East Asian systems (1,864), Ayurvedic and Indian traditional medicine (580), and both traditions (217). Point opacity is reduced for the dominant untagged class so that it does not visually obscure the tagged classes, and the cross-cultural class is drawn last. The subsample was drawn before the removal of 199 trivially small species, so approximately ten of the 50,000 members, and of those in Figures S6 and S7, are entries no longer present in the released corpus. Tagged compounds distribute across the macro-clusters rather than concentrating in any single region.

**Figure S6.**
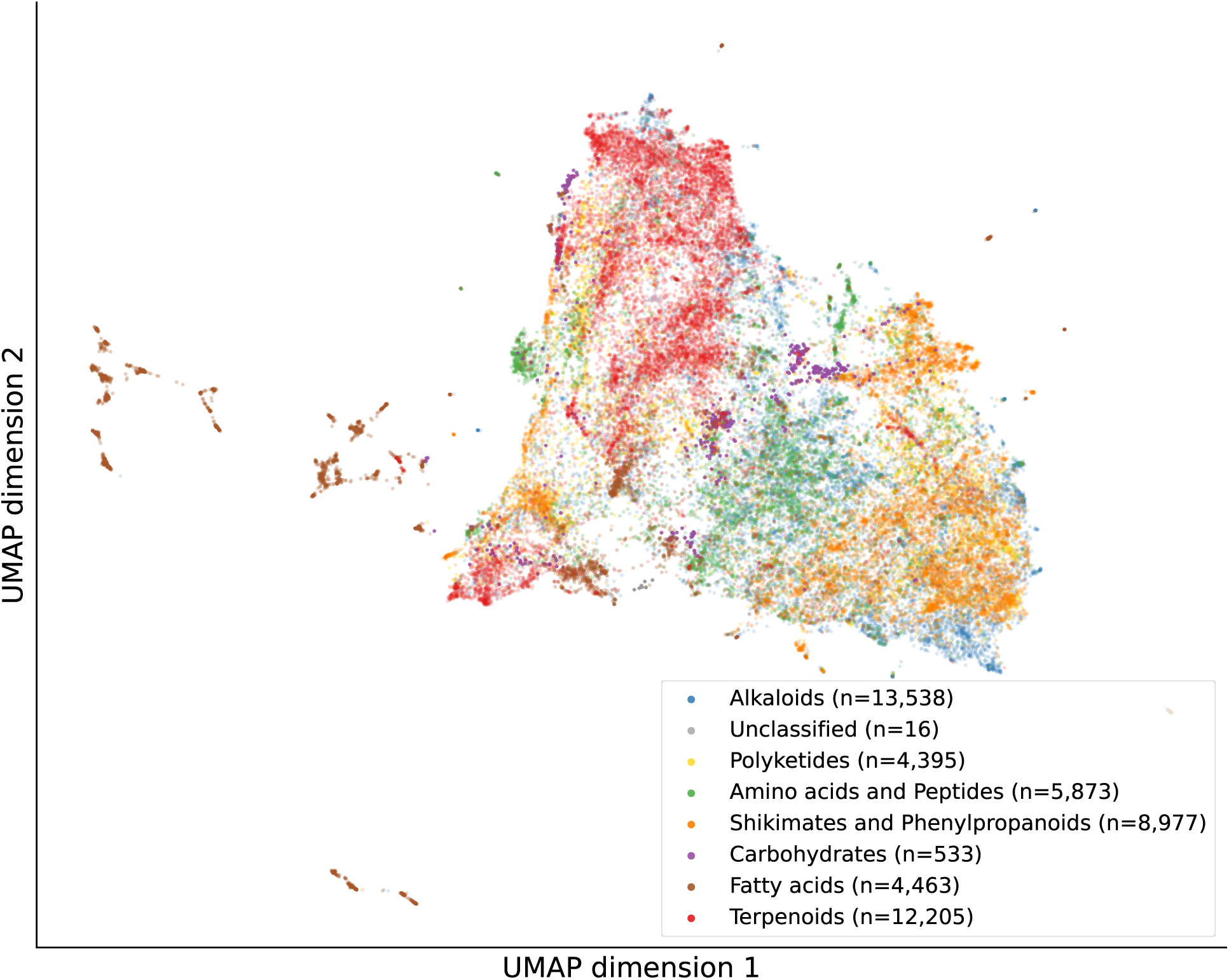
ChemBERTa embedding space colored by NPClassifier pathway. UMAP projection of the same mean-pooled 768-dimensional ChemBERTa embeddings (parameters as in Figure S5) across a uniform 50,000-compound subsample. Each compound is colored by effective pathway, resolved from the assigned class as in Figure S3: alkaloids (13,538), terpenoids (12,205), shikimates and phenylpropanoids (8,977), amino acids and peptides (5,873), fatty acids (4,463), polyketides (4,395), carbohydrates (533) and unclassified (16), comprising six subsample members with no ontology-derivable pathway and ten entries removed from the corpus after the embeddings were computed. The biosynthetic pathways occupy distinguishable regions of the embedding, which is consistent with the learned representation capturing pathway-relevant structure.

**Figure S7.**
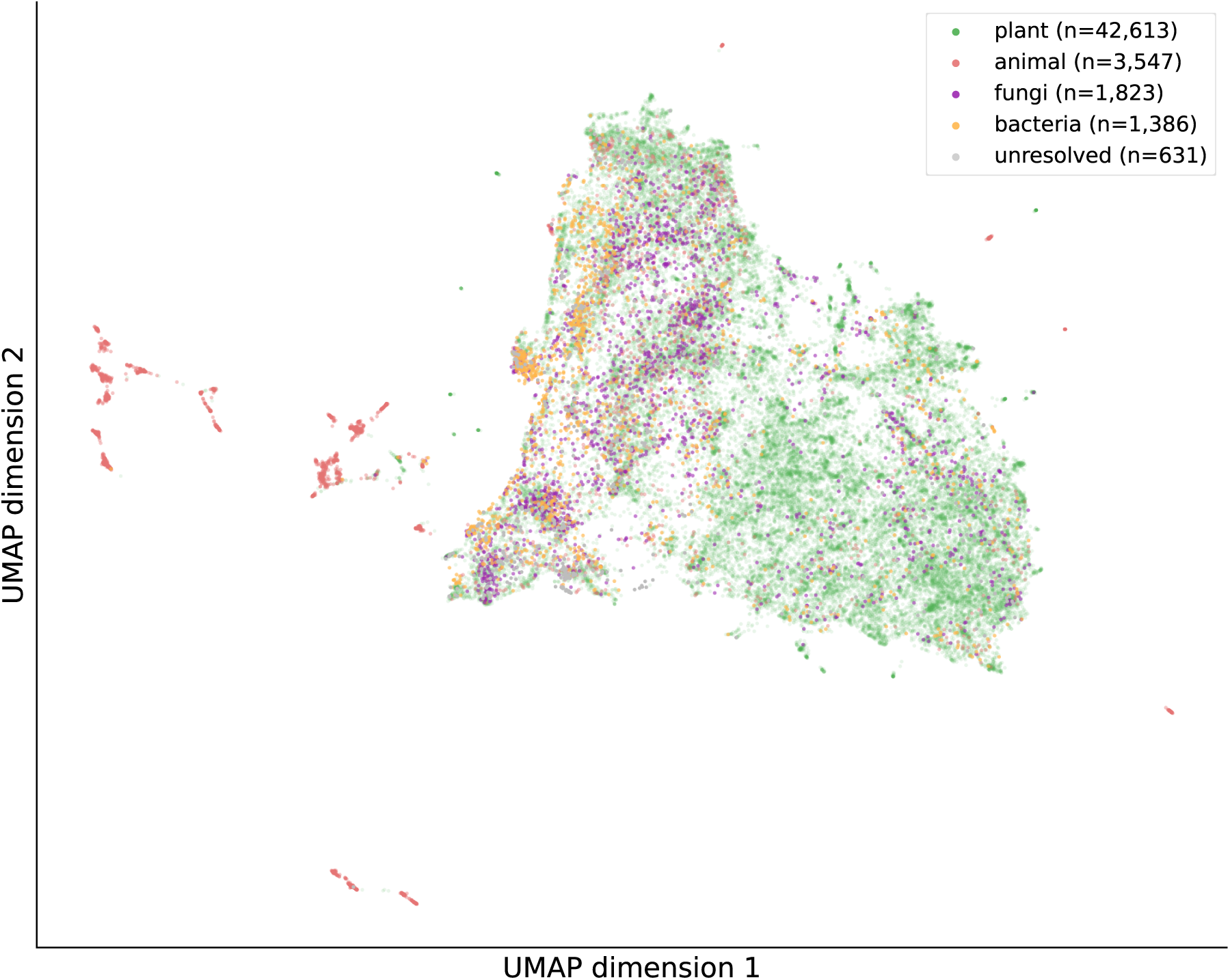
ChemBERTa embedding space colored by source-organism kingdom. UMAP projection of the same mean-pooled 768-dimensional ChemBERTa embeddings (parameters as in Figure S5) across a uniform 50,000-compound subsample as Figure S6: plant (42,613), animal (3,547), fungi (1,823), bacteria (1,386) and unresolved (631). Plant compounds dominate the visible distribution. Animal-, fungal-, and bacterial-derived compounds occupy distinguishable subregions, and unresolved-origin compounds distribute without strong localization. The same embedding therefore resolves three orthogonal axes of natural- product organization: traditional-medicine attestation (Figure S5), biosynthetic pathway (Figure S6), and taxonomic origin.

**Figure S8.**
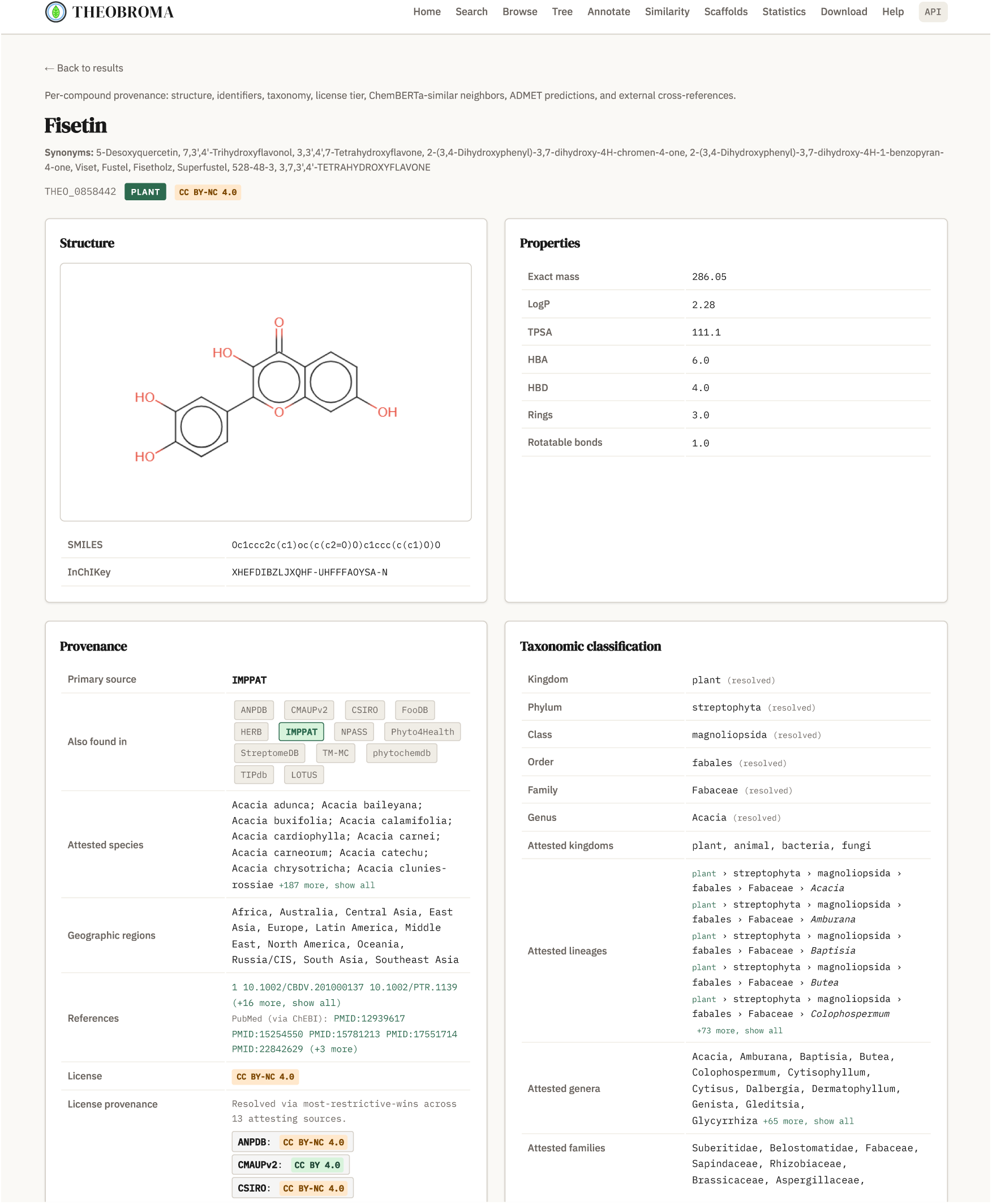
Compound result page for fisetin (THEO_0858442), part 1 of 3. The title block carries eleven PubChem synonyms including the trade names Fustel, Viset, Fisetholz and Superfustel, the CAS registry number 528-48-3, and systematic variants, alongside the primary identifiers (THEOBROMA ID, resolved kingdom, and per-compound license tier). Below it are the structure rendering with canonical isomeric SMILES and the full 27-character InChIKey, and the physicochemical descriptor panel (exact mass 286.05 Da, computed as RDKit ExactMolWt, LogP 2.28, TPSA 111.1, hydrogen-bond acceptors and donors, ring count, rotatable bonds). The provenance block lists all thirteen attesting databases, of which IMPPAT is the primary source, the attested species, the twelve mapped macro-regions, literature references via DOI and PubMed identifier, and the opening of the license provenance card continued in Figure S9. The taxonomic classification block gives the resolved lineage (kingdom plant, phylum Streptophyta, class Magnoliopsida, order Fabales, family Fabaceae, genus Acacia, each annotated “(resolved)”), the kingdom-restricted majority-voted output of the taxonomy resolver with the Streptophyta-phylum inference applied. Below it the attestation rows preserve the full per-source chain for audit: attested kingdoms, attested lineages ordered with the resolved family first, and attested genera and families. Long lists are truncated in the rendering with the remainder behind an inline expander and in full through the API. For this compound they hold 197 species, 78 lineages, 77 genera and 44 families.

**Figure S9.**
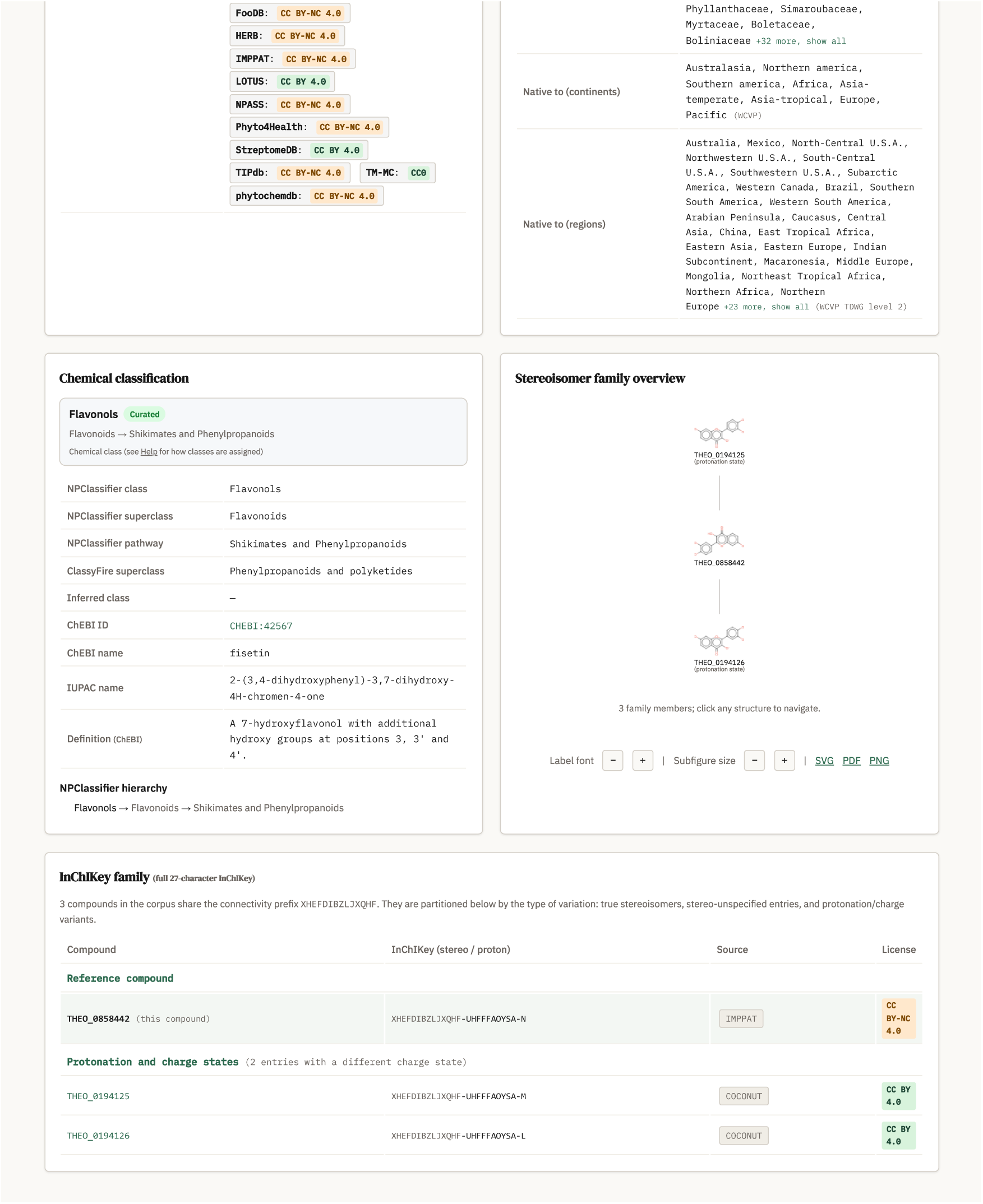
Compound result page for fisetin, part 2 of 3: the license provenance card continued from Figure S8, displaying the resolved-via-most-restrictive-wins outcome across 13 attesting sources (CC BY-NC 4.0 from ANPDB, CSIRO, FooDB, HERB, IMPPAT, NPASS, Phyto4Health, TIPdb and phytochemdb, CC BY 4.0 from CMAUPv2, LOTUS and StreptomeDB, CC0 from TM-MC), demonstrating the resolver where the attestation chain combines all three tiers and the most restrictive prevails. The WCVP native distribution follows at continent and TDWG level-2 region granularity. The chemical classification block carries the assigned class with its provenance badge (Flavonols, curated) above the NPClassifier hierarchy trace, the individual class, superclass and pathway fields, the ClassyFire (38) superclass, the inferred-class field left empty because fisetin is a curated-tier record, and the ChEBI cross-reference with IUPAC name and definition. The stereoisomer-family overview closes the page with an interactive radial display and the full-key family table, showing the three entries sharing the XHEFDIBZLJXQHF connectivity prefix: the reference record and two protonation-state variants contributed by COCONUT. The family spans license tiers, since the two COCONUT-only variants carry no non-commercial attestation and resolve to CC BY 4.0 where the reference record resolves to CC BY-NC 4.0. A license-filtered query therefore returns part of a connectivity family rather than all of it, which is a direct consequence of resolving licenses at compound rather than family granularity. The table partitions family members by type of variation. For fisetin only the protonation category is populated, since the flavonol scaffold has no stereocenters and no isotopologues are attested. The widget’s label-size, subfigure-size and export controls appear beneath it.

**Figure S10.**
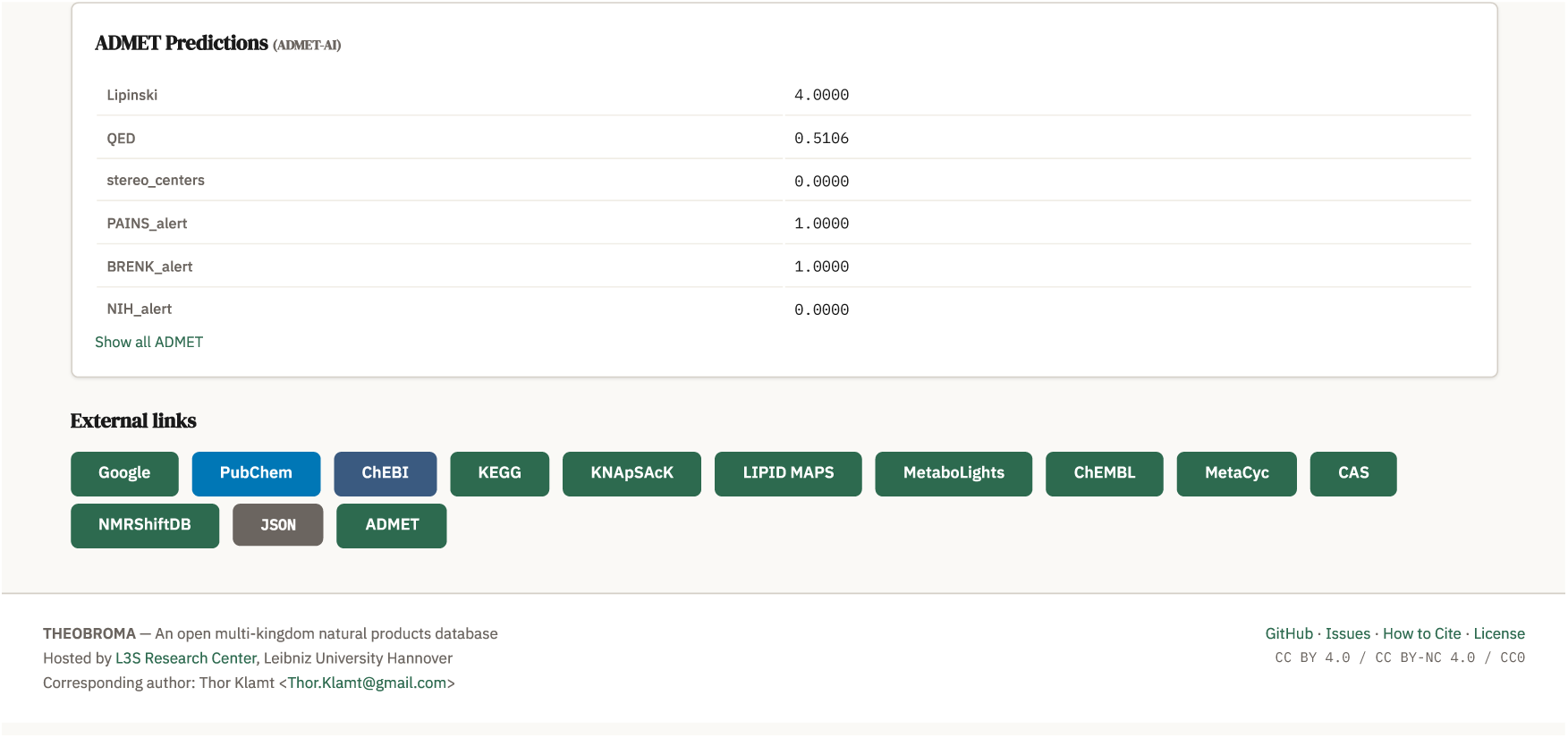
Compound result page for fisetin, part 3 of 3. The ADMET panel shows six default-visible items, which are RDKit-computed descriptors and structural-alert flags rather than ADMET-AI endpoints (Lipinski rules passed, QED drug-likeness, stereocenter count, and the PAINS, BRENK and NIH flags). The “Show all ADMET” expander reveals the full forty-seven-column ADMET annotation table on demand. The external cross-reference bar links to Google, PubChem, ChEBI, KEGG, KNApSAcK, LIPID MAPS, MetaboLights, ChEMBL, MetaCyc, CAS, NMRShiftDB, the JSON record and the ADMET detail view, with source-specific links rendered only where the compound is attested in that source. The page footer carries the THEOBROMA tagline, the L3S hosting attribution, project navigation links and the corpus-level license stripe spanning CC BY 4.0, CC BY-NC 4.0 and CC0.

**Figure S11.**
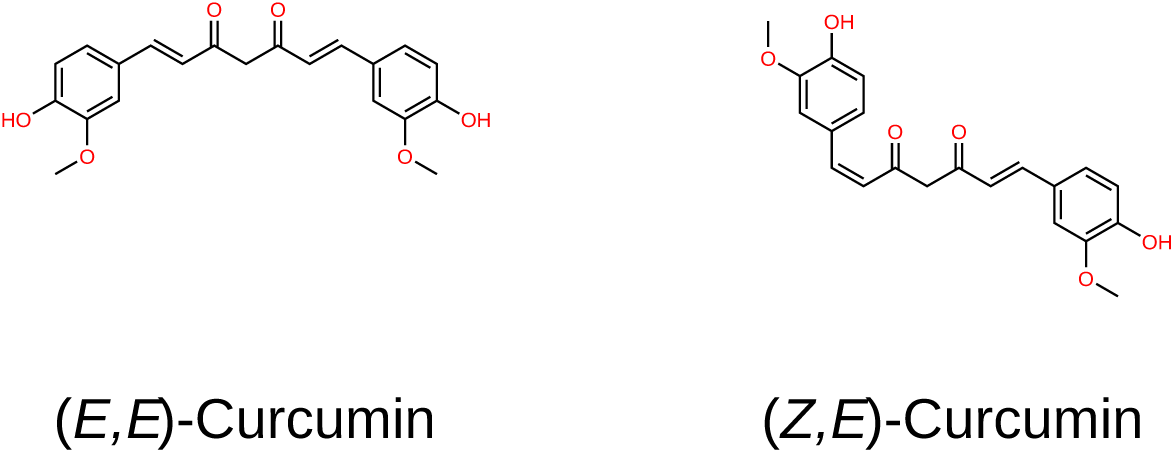
Complementing the example in the graphical abstract: curcumin stereoisomers preserved by full 27-character InChIKey deduplication. Left: (E,E)- curcumin, which does not measurably bind adenosine receptors A_1_AR or A_3_AR. Right: (Z,E)-curcumin, which binds A_1_AR with K_i_ = 306 nM and A_3_AR with K_i_ = 400 nM (29). Both are drawn as neutral parents. The corpus holds the (Z,E) configuration only in anionic form (THEO 0213373). They share the 14-character connectivity prefix, so databases deduplicating at that length collapse them and lose the binding distinction.

**Figure S12.**
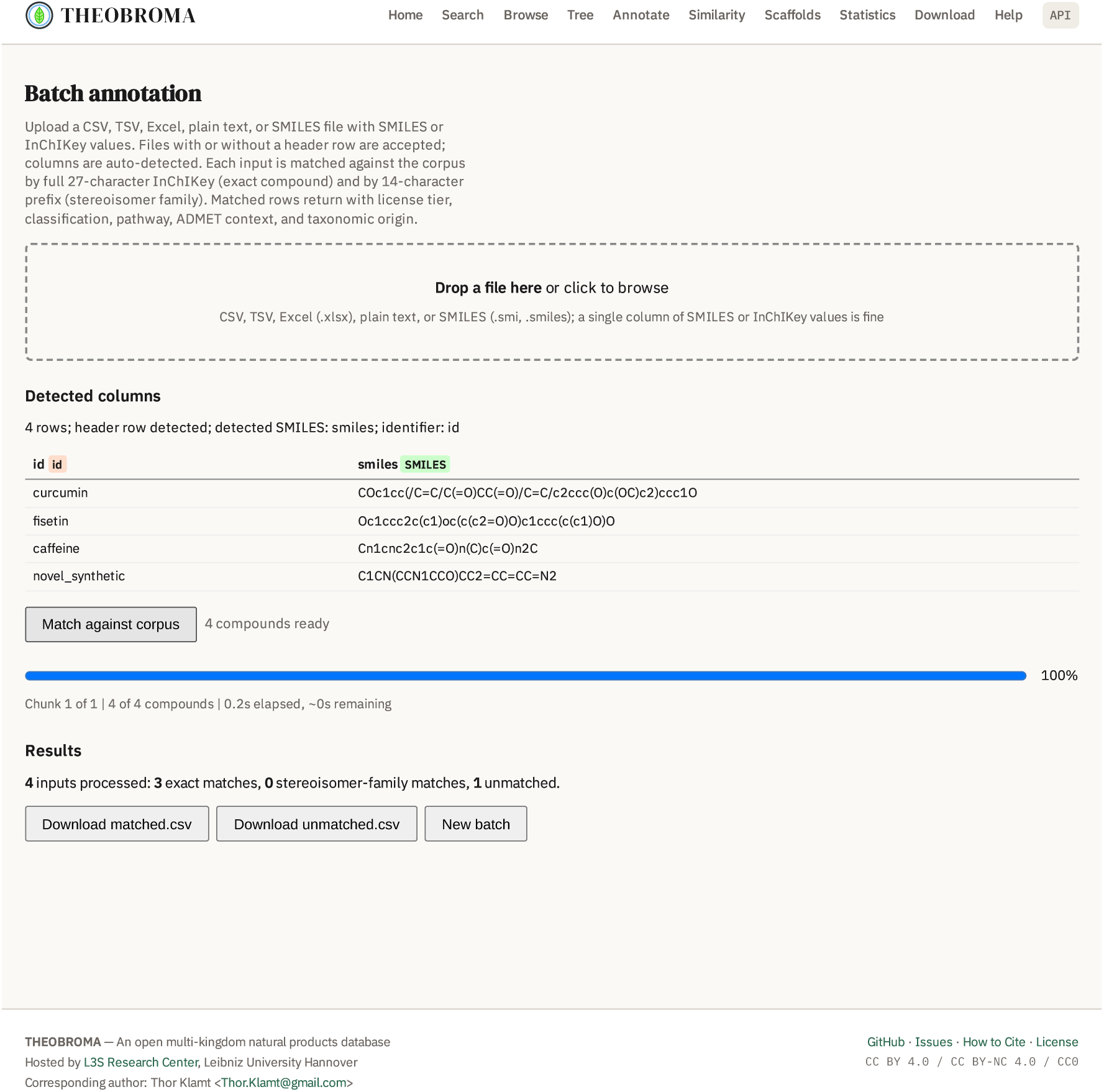
Batch annotation workflow demonstrated on a four-compound test input. The /annotate page identifies the SMILES column from its header, matched against a set of accepted names, and falls back to testing column values against structural patterns where no header matches. Processing proceeds through chunked client-side calls to /api/annotate, and a results summary distinguishes exact matches, stereoisomer-family matches, and unmatched compounds. Matched annotations and unmatched-input lists are independently downloadable as CSVs. Captured from the live deployment via headless-browser PDF render at viewport=1400×1600px, scale=0.7.

**Figure S13.**
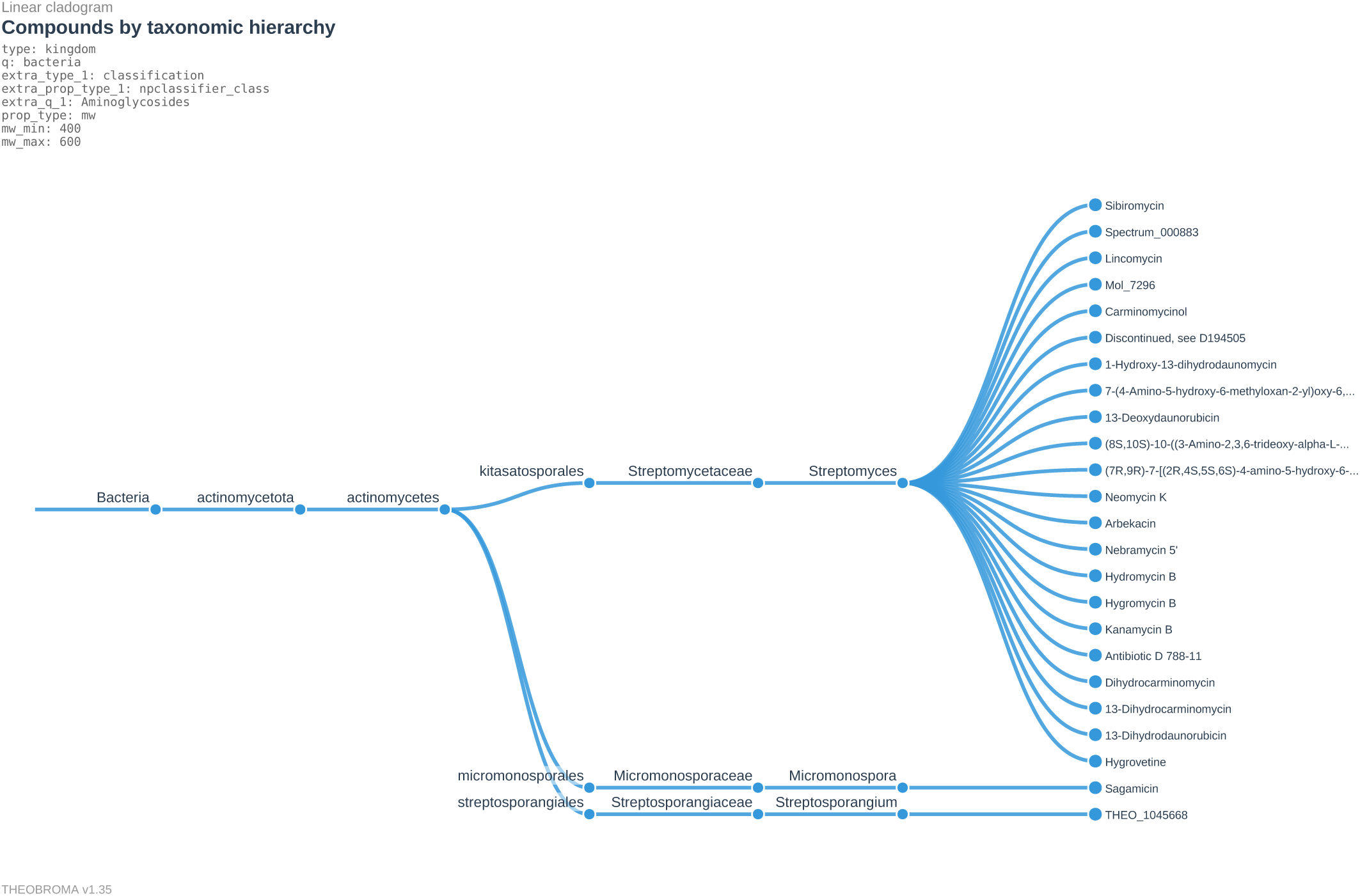
Compound-rank rendering of the THEOBROMA /tree cladogram for a bacterial query combining kingdom (bacteria), NPClassifier class (Aminoglycosides), and a molecular-weight band of 400 to 600 Da. The 30 returned compounds distribute across three actinomycete orders (Kitasatosporales, Micromonosporales, Streptosporangiales), three families, and three genera (*Streptomyces*, *Micromonospora*, *Streptosporangium*), and are shown as individual leaves beneath their genera, 24 of them rendered here, the remaining six having no resolved genus and therefore no branch to attach to, each terminal node carrying a compound name or THEOBROMA identifier. NPClassifier’s aminoglycoside class admits several amino-sugar-bearing scaffolds that are not aminoglycosides in the conventional sense, including anthracyclines, the lincosamide lincomycin and the pyrrolobenzodiazepine sibiromycin. The route hides compound leaves by default when a query returns more than one hundred compounds and shows them below that threshold, so the broader query of Figure 2 opens at genus rank while this one opens with compound leaves visible. The default can be overridden in the interface. The query composes a numeric property constraint with taxonomic and chemical filters, and renders a bacterial lineage rather than the plant lineage of Figure 2. Query terms, taxonomic hierarchy, and database version (v1.35) are baked into the PDF export.

**Figure S14.**
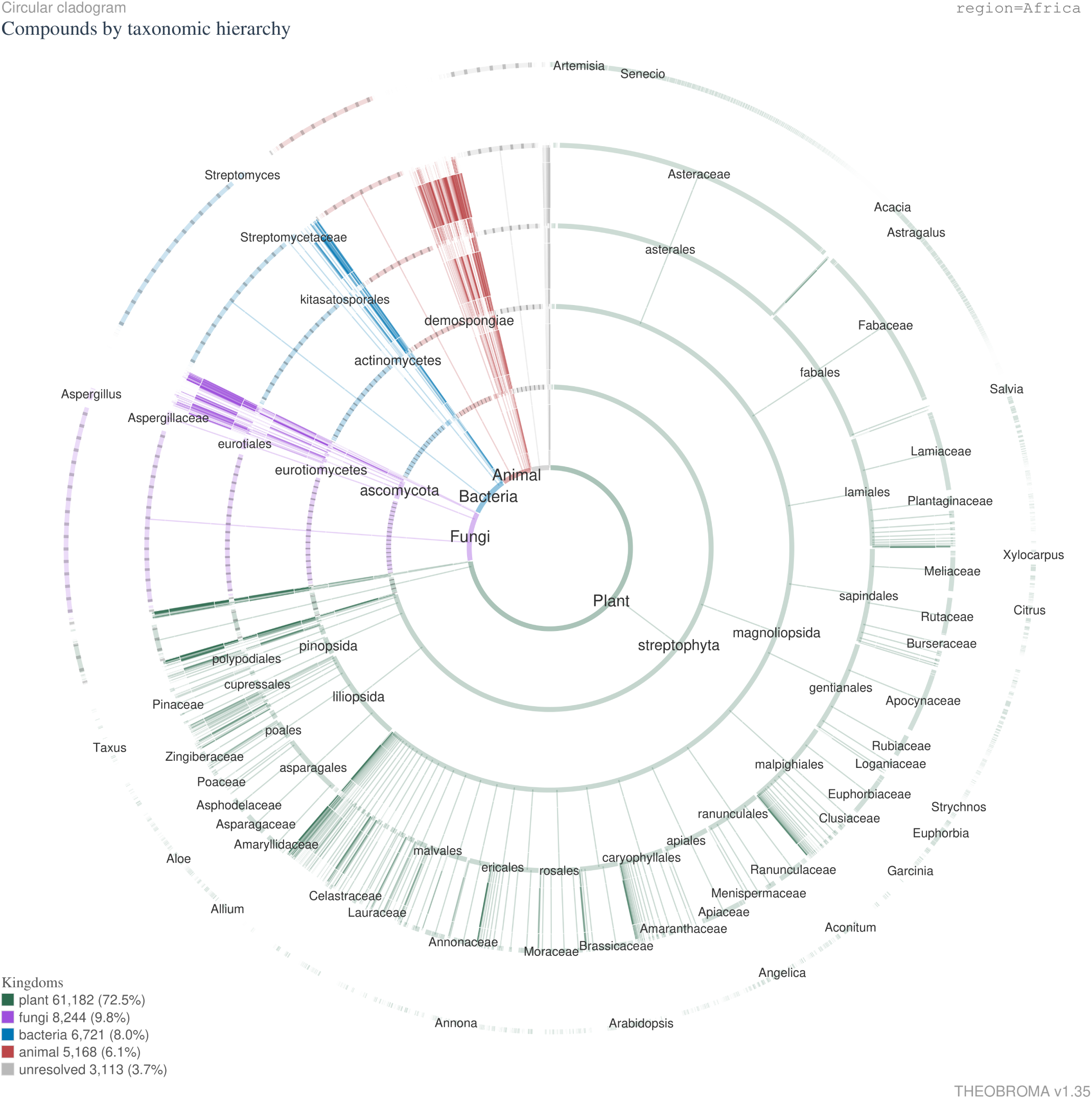
Radial rendering of the THEOBROMA /tree route for a single-axis region query (Africa), spanning compounds by taxonomic hierarchy across the kingdoms and phyla represented in that region. The 84,428 returned compounds distribute across plant (61,182, 72.5%), fungi (8,244, 9.8%), bacteria (6,721, 8.0%), animal (5,168, 6.1%) and unresolved (3,113, 3.7%). The radial layout distributes the taxonomic structure around the circumference, keeping the major clades distinct where a linear layout at this breadth would not. Query terms and database version (v1.35) are baked into the export.

**Table S1.** Kingdom distribution and secondary-kingdom cross-attestation after the v1.35 taxonomy rebuild and phylum reconciliation.

| Primary kingdom | Compounds |
| --- | --- |
| Plant | 964,360 |
| Animal | 80,485 |
| Fungi | 41,545 |
| Bacteria | 32,749 |
| Unresolved | 13,666 |
| Secondary-kingdom attestations | Compounds |
| One secondary kingdom | 8,868 |
| Two secondary kingdoms | 4,923 |
| Three secondary kingdoms | 229 |
| Total with secondary attestation | 14,020 |

**Table S2.** XGBoost hyperparameter search space and selected values for the deployed classifier (Optuna, 75 requested trials completing at 78 because workers had further trials in flight when the target was reached under four-worker parallel execution, 50 random startup trials followed by TPE-guided search, objective 5-fold cross-validated macro-F1, best trial validation macro-F1 0.812, the Optuna objective value, against the 0.809 five-fold cross- validated mean reported in the main text for the deployed configuration). The feature vector is 2,471-dimensional: 2,048-bit Morgan fingerprints, 167 MACCS keys, and a 256-dimensional PCA projection of the ChemBERTa embedding, over 501 classes. Every selected continuous value lies in the interior of its search range. The categorical weight mode selected none, an endpoint of its three-value set, which the cross-validated objective preferred over sqrt and full. The deployed model is trained to 467 boosting rounds under early stopping rather than to a fixed round count. The unweighted configuration was selected under the cross-validated macro-F1 objective. On a correctly derived pathway the model tier assigns Alkaloids at 40.6% against 25.4% for the curated tier, and class-balanced retraining remains the documented route to reducing that margin.

| Hyperparameter | Search range | Selected |
| --- | --- | --- |
| `n_rounds` | $\leq 800$ (early stop) | 467 |
| `max_depth` | 4–14 | 9 |
| `learning_rate` | 0.05–0.5 (log) | 0.0888 |
| `subsample` | 0.5–1.0 | 0.876 |
| `colsample_bytree` | 0.1–1.0 | 0.692 |
| `min_child_weight` | 1–30 | 3 |
| `lambda (L2)` | 0.01–50 (log) | 1.412 |
| `alpha (L1)` | 0.001–20 (log) | 0.00598 |
| `gamma` | 0–5 | 0.184 |
| `weight_mode` | none / sqrt / full | none |

**Table S3.**
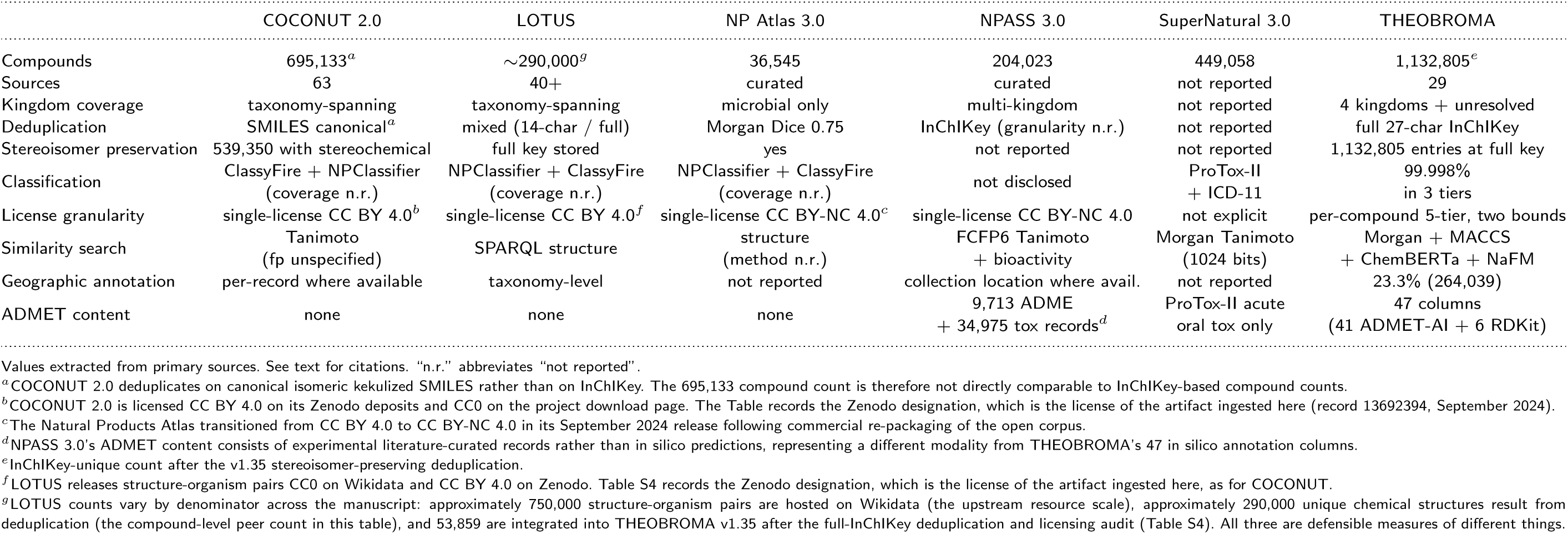
Full comparison of THEOBROMA against other open natural-product aggregators. The main text reproduces the three differentiator axes only. Cells marked “not reported” indicate

**Table S4.**
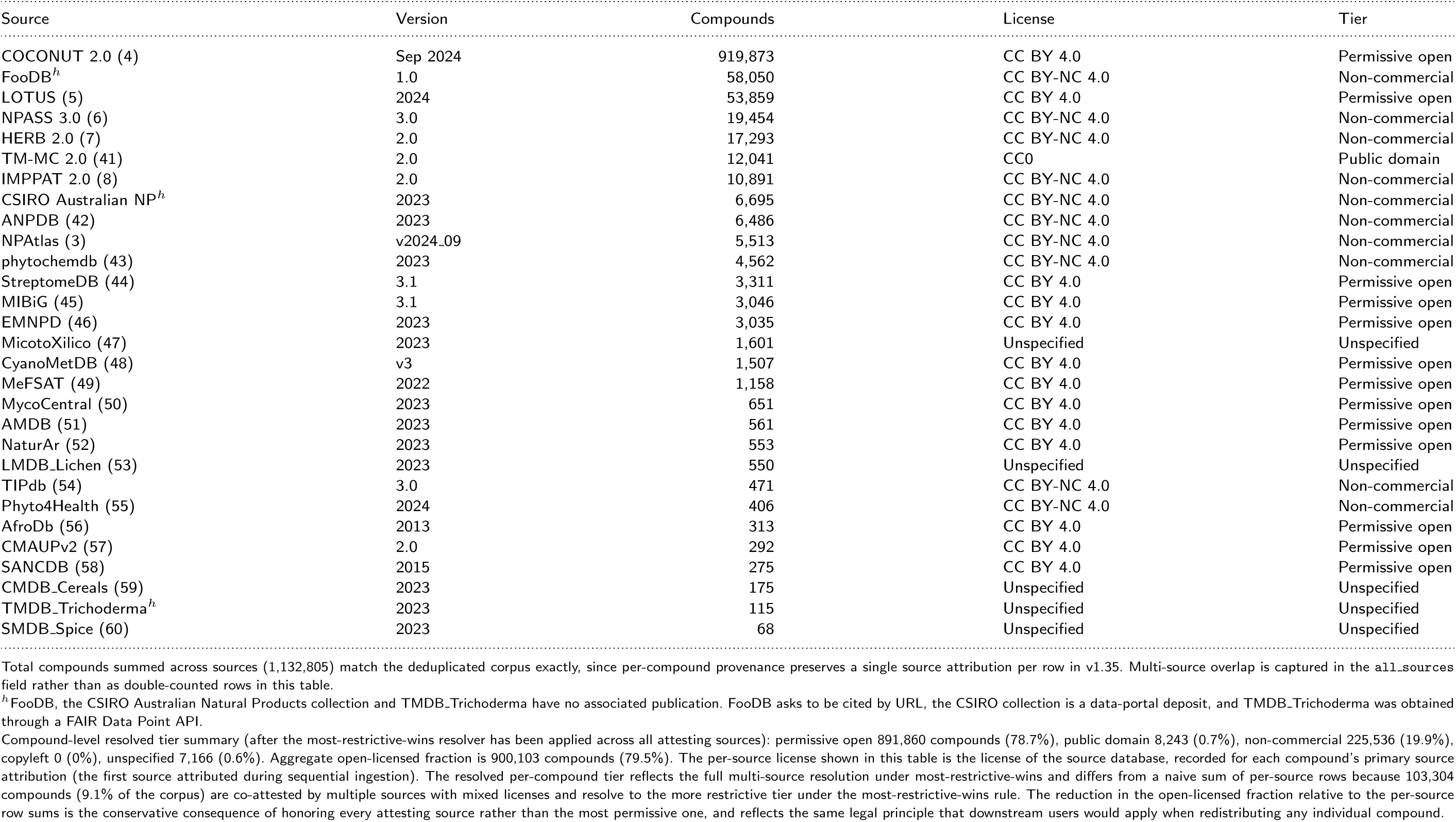
Per-source license audit across the 29 integrated sources. Compound counts reflect post-deduplication integration and primary attribution. The Version column records the upstream release identifier where the source declares one and the download snapshot year otherwise, as recorded in the reproducibility manifest. License tiers follow the five-tier taxonomy of Supplementary S3.

